# Unique structural features of mammalian NEIL2 DNA glycosylase prime its activity for diverse DNA substrates and environments

**DOI:** 10.1101/2020.02.28.970129

**Authors:** Brian E. Eckenroth, Vy Cao, April M. Averill, Julie A. Dragon, Sylvie Doublié

**Affiliations:** Department of Microbiology and Molecular Genetics, University of Vermont, Stafford Hall, 95 Carrigan Drive, Burlington, Vermont 05405, United States

**Keywords:** DNA damage, DNA repair, NEIL2 glycosylase, SAXS, crystallography

## Abstract

Oxidative damage on DNA arising from both endogenous and exogenous sources can result in base modifications that promote errors in replication as well as generate sites of base loss (abasic sites) that present unique challenges to maintaining genomic integrity. These lesions are excised by DNA glycosylases in the first step of the base excision repair pathway. Here we present the first crystal structure of a NEIL2 glycosylase, an enzyme active on cytosine oxidation products and abasic sites. The structure reveals an unusual “open” conformation not seen in NEIL1 or NEIL3 orthologs. NEIL2 is predicted to adopt a “closed” conformation when bound to its substrate. Combined crystallographic and solution scattering studies show the enzyme to be conformationally dynamic in a manner distinct among the NEIL glycosylases and provide insight into the unique substrate preference of the enzyme. Additionally, we characterized three cancer variants of human NEIL2, namely S140N, G230W, and G303R.

## INTRODUCTION

Sites of oxidative damage on DNA arise from numerous sources including environmental agents, endogenously produced oxygen species from normal cellular respiration, and other vital processes like demethylation, which produce intermediates including abasic sites. Improperly managed or repaired sites can result in polymerase stalling during replication, and ultimately mutagenesis and cancer. Mammals employ a battery of enzymes aimed at repair of DNA damage for the faithful replication of the genome and/or transcript interpretation. One such enzyme, Nei-like 2 (NEIL2), belongs to the Fpg/Nei family of DNA glycosylases (**Figure 1**) of the base excision repair pathway (BER). This bifunctional enzyme is capable of excising an oxidized base (glycosylase activity) as well as cleaving the DNA backbone (lyase activity) with the ability to function in an apurinic endonuclease (APE)-independent manner (1). NEIL2 displays activity on both single- and double-stranded DNA with a preference for bubble DNA structures, typically selecting oxidation products of cytosine(2). Proteins of the Fpg/Nei family display remarkable structural conservation of the two-domain architecture while being highly divergent at the sequence level, particularly within the predominantly β-strand-rich N-terminal domain. The C-terminal domain contains the conserved helix-two-turn-helix (H2TH) and zinc or zincless finger(3) DNA binding motifs and is reasonably conserved within the family.

**Figure 1.**
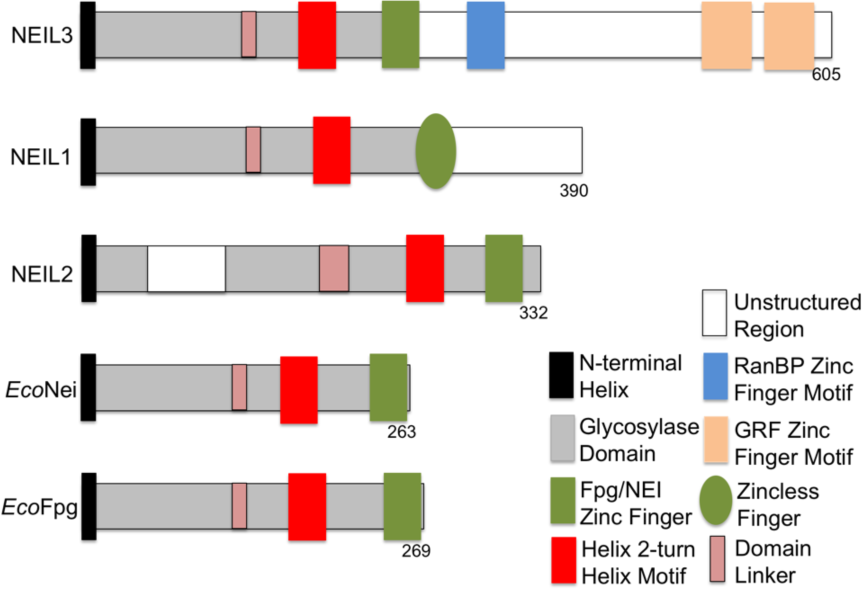
Diagram of Fpg/Nei glycosylases highlighting their structural features. NEIL enzymes are substantially larger than their bacterial counterparts. The eukaryotic enzymes harbor disordered regions. NEIL2 is unique in that the flexible region is internal. Figure adapted from(47).

Accumulating evidence suggests that NEIL2 activity likely fulfills a context-dependent function rather than simple global DNA damage surveillance(4). Studies using NEIL2 knock-down mouse models have produced mixed results with regard to the accumulation of oxidative DNA damage(5, 6), with biochemical and cell-based data indicating a preference for repair of damage in transcriptionally active regions of the DNA(7). This is in line with the correlation to transcription associated repair via the observed interaction of NEIL2 with Cockayne Syndrome Protein B(8). While a direct linkage to a cancer phenotype is yet to be established for NEIL2, studies have identified elevated risk associations to carriers of BRCA2 mutations – identifying NEIL2 as a risk modifier. Single nucleotide polymorphism in the NEIL2 gene evaluated from patients in the CIMBA consortium showed significant association between NEIL2 and BRCA2 mutation carriers and breast cancer risk(9) while further analysis suggested that the elevated production of NEIL2 resulting from this SNP correlated with elevated oxidative DNA damage for the BRCA2 mutation carriers(10). Recent developments also point to alternative functions of the NEIL glycosylases beyond DNA repair via their contributions to the demethylation pathway of 5-methyl cytosine(11) (12).

While some hereditary-based cancers follow a predictable paradigm based on mutations to a specific gene or set of genes, the majority of cancers are considerably more complex and likely arise from an ensemble of mutations to an array of gene products modulated by the basal genomic variability within individual patients. Identifying all contributing factors and establishing their contextual roles are essential to understanding the development and progression of cancers as well as selection of the most effective therapeutic protocol. Herein we report the first X-ray crystal structure of any Neil2 enzyme, in concert with solution-based studies. The protein adopts a unique open orientation in the absence of DNA substrate, predicting a necessary large conformational change to assemble a catalytically competent complex. The results provide insight into the substrate diversity of NEIL2 and shed light on a unique protein appendage that would allow interactions with multiple protein partners without impeding enzyme function. *In vitro* activity of cancer-associated variants of NEIL2 suggests that structural perturbations impact the overall enzymatic activity with significant downstream implications for BER.

## MATERIALS and METHODS

### Protein Expression and Purification

Both full-length and truncated gray short tailed opossum *Monodelphis domestica* Neil2 (MdoNEIL2) sequences were synthesized and codon-optimized for expression in *E. coli* by GenScript and subcloned into a pET30 vector (Novagen) using NdeI and XhoI restriction sites, then transformed into BL21(DE3) plated onto LB-agar under kanamycin selection. Positive colonies were used for bulk expression using Terrific Broth (TB) supplemented with 4% (v/v) glycerol and 10 μM zinc sulfate. Culture was grown to an OD_600_ of 1.0, induced with 500 μM IPTG, and expressed for 6 hours at room temperature. Cells were harvested then lysed in buffer containing 50 mM sodium phosphate pH 8, 300 mM NaCl, 10% (v/v) glycerol, 10 mM imidazole, 2 mM beta-mercaptoethanol, 0.01% (v/v) NP-40 and 1 mM PMSF and purified by nickel affinity resin (Thermo Fisher Scientific) with elution in buffer containing 400 mM imidazole. The eluant was exchanged to 25 mM Hepes pH 7.5, 100 mM NaCl, 10% (v/v) glycerol and 1 mM DTT prior to ion exchange chromatography using a Fast Flow SP column (GE Healthcare). Elution from the SPFF column was performed with a 100 mM to 1M gradient over 20 column volumes. Concentrated aliquots were stored in 50 mM HEPES pH 7.5, 300 mM NaCl, 1 mM DTT and 10% glycerol and flash frozen in liquid nitrogen for storage at −80°C. Expression for selenomethionine incorporation substituted the TB media with minimal media as described in(13). For phasing verification three sites were mutated from Leu to Met (7, 210 and 284) to increase the number of heavy atom sites described below. Upon structure solution of the full-length protein, a construct lacking the internal disordered region was engineered, replacing the [F68-N127] segment with a short flexible linker of sequence GSGSG. The loop deletion construct for MdoNEIL2 was expressed and purified as described above. Purification of human NEIL2 (HsaNEIL2) followed the same protocol as MdoNEIL2, with the exception of the expression step: The pET30-HsaNEIL2 construct is not codon optimized for *E. coli* expression and thus required Rosetta2 DE3 pLysS cells (Novagen) for expression.

### Glycosylase Activity Assays

Initial activity for NEIL2 was screened using a cyanoborohydride trapping assay. Reactions containing 17 μM enzyme and 43 μM DNA substrate were incubated at 25°C for 30 minutes in 10 mM Tris-HCl pH 8.0, 50 mM NaCl, 1 mM DTT, 1 mM MgCl_2_ in the presence of 100 mM sodium cyanoborohydride. Reactions were quenched by the addition of 100 mM Tris and run on a denaturing 12% SDS-PAGE gel. Kinetic parameters were determined using ^32^P labeled DNA oligos and performed using single-turnover and multiple-turnover conditions. The 35-mer oligonucleotides with damage-containing strand sequence of 5′-TGTCAATAGCAAG(X)GGAGAAG TCAATCGTGAGTCT-3′, where X was Tg, 5-OHU, 5,6-DHU were purchased from Midland Certified Reagent Co. (Midland, TX), purified by urea PAGE and labeled using T4 polynucleotide kinase in the presence of γ-^32^P as previously described(14). Single-turnover assays were performed using 25 nM DNA and 25-800 nM enzyme for 30 minutes at 37°C in 10 mM Tris-HCl pH 8.0, 75 mM NaCl, 100 ug/mL BSA and 1 mM DTT. Maximum activity fraction under the assay conditions for the most active enzymes could be estimated using fits to standard single exponential curves however, assays with lower activity fit poorly. All assay titration curves are provided in **Supplemental Figure 3**. Final comparison of maximal activity was performed using the mean activity at the highest enzyme concentration and shown in **Figure 3**.

**Figure 2.**
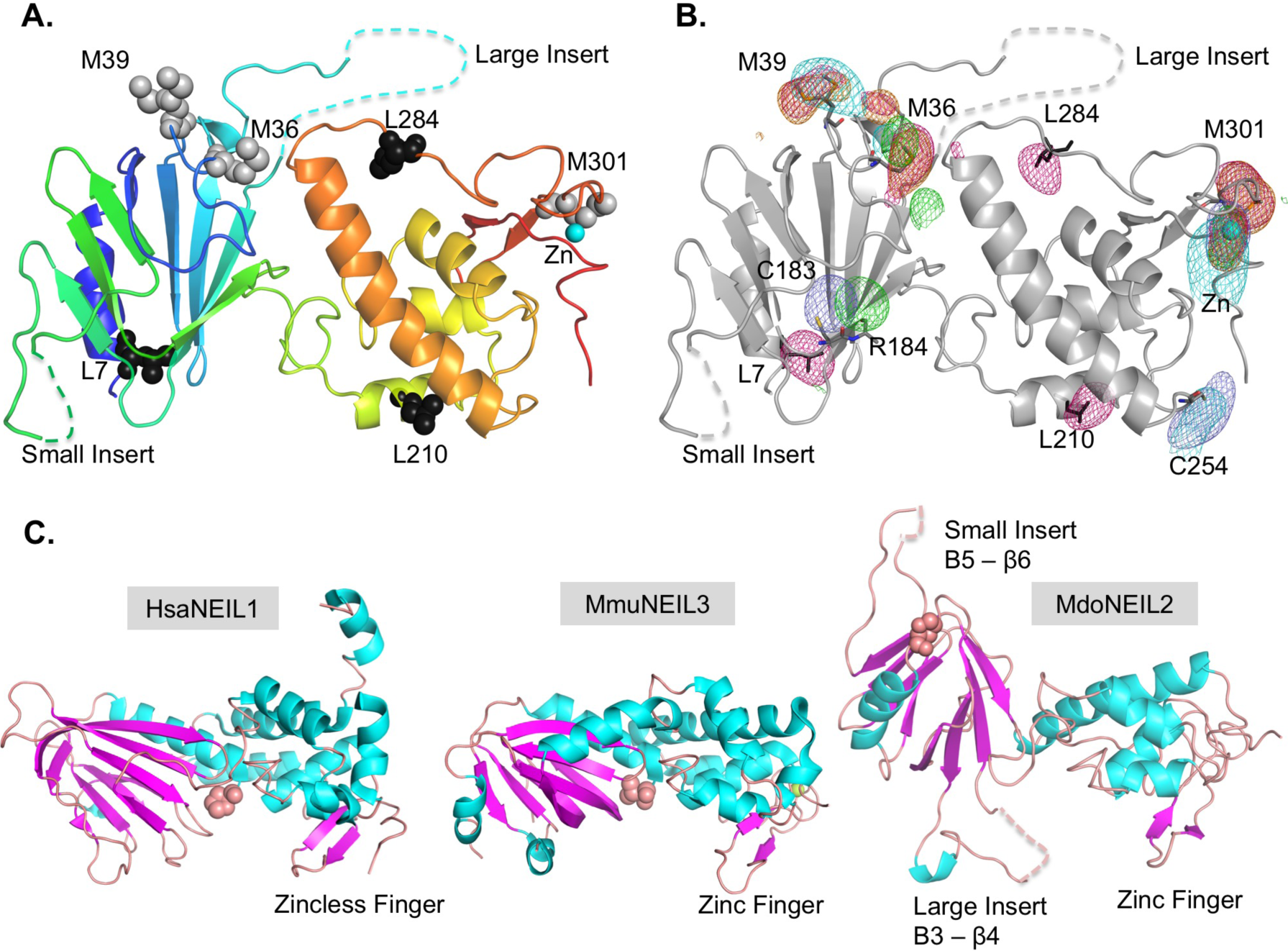
Structure of MdoNEIL2 and comparison to other glycosylases. A.) Cartoon representation of NEIL2 structure from N-terminus (blue) to C-terminus (red) with wild-type methionines highlighted with grey spheres and methionines engineered for additional phase verification shown in black. B.) Cartoon representation of NEIL 2 (gray) and overlaid anomalous difference Fourier maps contoured at 3σ for the WT SeMet (orange), L-M SeMet (purple), sodium iodide soaked (green), KAu(CN)_2_ soaked (blue), K_2_PtCl_4_ (cyan). C.) Comparison of MdoNEIL2 to human NEIL1 (HsaNEIL1; PDB ID code: 1TDH) (3) and mouse NEIL3 (MmuNEIL3; PDB ID code: 3W0F)(34) displayed with the C-terminal domain in equivalent orientations and demonstrating the unique inter-domain orientation of NEIL2. The N-terminal active site residue is shown in pink, in space-fill mode.

**Figure 3.**
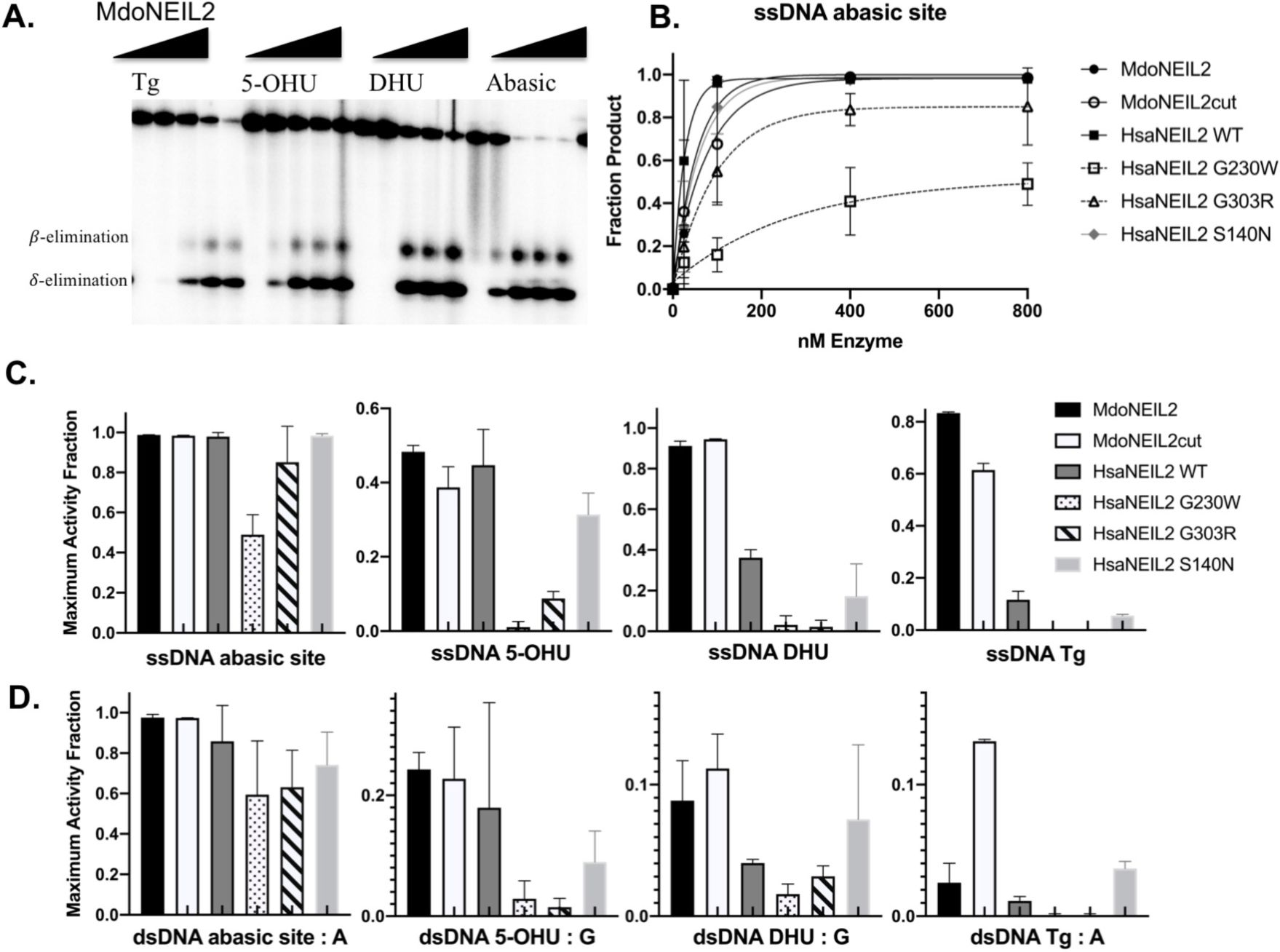
Activity of MdoNEIL2 towards DNA lesions. A.) Glycosylase activity of MdoNEIL2 under single-turnover conditions for ^32^P-labelled ssDNA substrates run on urea-PAGE. Assays were performed using 25 nM DNA substrate and enzyme concentration titrated from 0 to 800 nM. B.) Shown is the activity titration for single stranded abasic site substrate with data fit to a single exponential. C.) Shown is the summary of the maximum activity fraction for all single stranded substrates and enzyme variants of MdoNEIL2 and HsaNEIL2. D.) Shown is the summary of the maximum activity fraction for all double-stranded substrates

### Crystallization and Data Collection of opN2

Frozen aliquots of full-length MdoNEIL2 were thawed and exchanged into buffer containing 50 mM HEPES pH 7.5, 100 mM NaCl and 1 mM TCEP. Crystallization optimization was performed by hanging drop vapor diffusion after high-throughput screening identified suitable starting conditions (PEG-Ion HT, condition C11; Hampton Research). For the initial matrix screening, the protein was retained in the storage buffer containing 10% (v/v) glycerol. Crystallization hits were amorphous quasi-crystals with subsequent experimentation showing removal of the glycerol to be essential for producing properly formed crystals. Protein concentration was optimal between 1.5 and 3 mg/ml. Optimized conditions contained 40 mM HEPES pH 7.5, 1 mM TCEP, 50 mM sodium succinate, 1% propylene glycol and 12-16% PEG 3350. Cryoprotection was achieved by increasing the PEG concentration to 20%, with the inclusion of 30% glucose. Crystals diffracted to 2.7 Å for the native full-length opN2, 2.5 Å for the selenomethionyl version, and 3.1 Å for the selenomethionyl L7/210/284M triple mutant. Heavy atom soaks used native crystals and were performed in the cryoprotection stabilizing reagent for 2 hours at room temperature in the presence of 200 mM NaI, 5 mM KAu(CN)_2_, or 5 mM K_2_PtCl_4_. Crystals were screened for diffraction quality on a home source (Bruker D8 Quest). High-resolution data were acquired at the Advanced Photon Source GM/CA Sector 23 on either a Mar300 CCD or Pilatus 6M detector. Data for each heavy atom were collected at the peak wavelength (Table 1), with the WT selenomethionine also collected at a high-energy remote wavelength. The truncated form of MdoNEIL2 crystallized in similar conditions to that of full-length, with the distinction that crystals were consistently smaller and diffracted poorly.

**Table 1.** Crystallographic data collection statistics used for structure determination of MdoNEIL2. The data for phase determination were scaled anisotropically with ellipsoidal truncation using STARANISO; anisotropy resolution limits provided along a* and c* for the datasets are indicated. Data sets for validation were processed utilizing Proteum3. Final model refinement was performed using spherically truncated data shown in Table 2.

|  | WT Native | WT SeMet Peak | WT SeMet Remote | WT Zn Edge | WT Iodide | WT Platinum Peak | WT Gold Peak | L7,210, 284M SeMet Peak |
| --- | --- | --- | --- | --- | --- | --- | --- | --- |
| Number of crystals | 2 | 2 | 2 | 1 | 1 | 1 | 1 | 1 |
| Wavelength Å | 1.033 | 0.979 | 0.870 | 1.28 | 1.77 | 1.072 | 1.039 | 0.979 |
| Space Group | P3(2) | P3(2) | P3(2) | P3(2) | P3(2) | P3(2) | P3(2) | P3(2) |
| Cell a=b, c | 68.3, 149.2 | 67.9, 149.1 | 67.9, 149.3 | 68.2, 148.2 | 67.8, 147.3 | 68.9, 151.6 | 68.0, 148.5 | 67.7, 149.1 |
| Resolution Å (High resolution Å) | 58.9-2.63 (2.79-2.63)<sup>a</sup> | 40-2.54 (2.68-2.54) | 34.02-2.75 (2.94-2.75) | 59-2.95 (3.05-2.95) | 58.9-3.07 (3.36-3.07) | 39-3.89 (4.03-3.89) | 58.9-2.79 (2.98-2.79) | 38-3.1 (3.21-3.1) |
| Resolution $a^*$ Å | 2.91 | 2.86 | 2.96 | | 3.32 | | 3.22 | |
| Resolution $c^*$ Å | 2.63 | 2.54 | 2.75 | | 3.07 | | 2.79 | |
| Reflections | 22928 | 25470 | 20072 | 16233 | 14323 | 7451 | 18570 | 13806 |
| Completeness % | 100 (100) | 100 (99.9) | 99.8 (100) | 99.6 (96.4) | 99.1 (100) | 99.6 (99.0) | 96.2 (99.7) | 99.6 (96.8) |
| Multiplicity | 16.2 (9.7) | 11.0 (9.0) | 7.3 (7.2) | 5.8 (5.6) | 9.1 (7.5) | 3.6 (3.4) | 5.7 (5.9) | 5.4 (5.4) |
| Wilson B | 63.2 | 64.2 |  |  |  |  |  |  |
| I/SigmaI | 14.1 (1.4) | 18.7 (0.9) | 17.6 (1.1) | 7.8 (0.9) | 13.1 (1.2) | 6.5 (1.4) | 9.3 (0.8) | 8.5 (1.2) |
| Rmeas % | 18.4 (575) | 13.0 (376) | 10.2 (197) | 23.3 (369) | 25.3 (245) | 15.4 (91.8) | 16.2 (288) | 13.9 (143) |
| Rpim % | 4.4 (155) | 3.9 (121) | 3.8 (72.8) | 9.7 (155) | 11.8 (126) | 8.2 (49.8) | 6.9 (117) | 6.0 (60.8) |
| CC ½ | 99.9 (55.3) | 99.8 (46.8) | 99.7 (60.1) | 99.5 (33.0) | 99.5 (53.1) | 99.3 (29.2) | 99.6 (51.1) | 99.7 (48.1) |
| D-Anom 1/2 sets |  |  |  |  |  |  |  |  |
| Overall |  | 0.53 |  |  | 0.1 |  | 0.15 |  |
| Inner shell |  | 0.61 |  |  | 0.24 |  | 0.39 |  |
| Outer shell |  | 0.01 |  |  | 0.02 |  | 0 |  |
| Anomalous cutoff Å |  | 3.7 |  |  | 5.4 |  | 5.7 |  |
| Cross R<sub>factor</sub> F<sub>obs</sub> % | --- | 12.0 | 13.7 | 23.2 | 23.2 | 76.0 | 26.4 | 22.5 |
| Phasing Power |  |  |  |  |  |  |  |  |
| Isomorphous |  | 1.30 |  |  | 0.47 |  | 0.51 |  |
| Anomalous |  | 0.77 |  |  | 0.29 |  | 0.48 |  |
| Heavy Atom Sites |  | 10 |  |  | 6 |  | 6 |  |
| CCall/CCweak |  | 45.2/27.0 |  |  | 34.7/15.3 |  | 50.0/30.6 |  |
| CFOM |  | 72.2 |  |  | 50.0 |  | 80.6 |  |
| Correlation |  | 0.35 |  |  | 0.28 |  | 0.46 |  |
| Phasing R-cullis |  |  |  |  |  |  |  |  |
| Isomorphous |  | 0.42 |  |  | 0.84 |  | 0.92 |  |
| Anomalous |  | 0.87 |  |  | 0.98 |  | 0.96 |  |
| Phasing FOM |  | 0.312 |  |  |  |  |  |  |
| (E**2 corr)/contrast |  |  |  |  |  |  |  |  |
| Density Modif. |  | 0.45 |  |  |  |  |  |  |
| Solvent Flat. |  | 3.1 |  |  |  |  |  |  |
| Initial Model |  |  |  |  |  |  |  |  |
| R/R<sub>free</sub> % |  | 21.6/39.3 |  |  |  |  |  |  |

<sup>a</sup>Numbers in parentheses denote high-resolution bin.

### *Structure determination of* MdoNEIL2

Diffraction data were processed with XDS(15) and scaled using STARANISO(16), with the exception of the data sets for the selenomethionyl L7/210/284M variant, Platinum derivative, and the truncated form of MdoNEIL2, which were processed with Proteum3 (Bruker AXS). Initial selenomethionine sites for the full-length construct along with anomalous scattering from the zinc atom were identified using ShelxD(17) as incorporated into AutoSHARP(18). The full-length protein sequence contains 336 residues and 4 methionines. Sites were refined using ShelxE and phasing performed using AutoSHARP followed by improved density modification using Solve/Resolve within Phenix(19). Cross comparison between datasets was performed using Scaleit after data merging using CAD. An initial solution was achieved using space group P3_2_12, as suggested by Laue group interpretation and self-rotation function analysis. Model building and refinement stalled with R-factors greater than 30% and elevated B-factors. Detailed analysis of anomalous reflection correlations identified the lower symmetry space group P3_2_ as a more probable solution. A final phased solution using AutoSHARP was achieved using the WT selenomethionine, iodide and gold derivatives using STARANISO ellipsoidal truncated data. The L7/210/284M selenomethionyl variant, WT platinum, and a native crystal collected at the peak wavelength for zinc were used for validation (**Table 1**). Model building was performed in Coot(20) and final refinement performed with Phenix using the spherically truncated data processed by either Proteum3 for the single highest resolution crystal of the WT selenomethionine or aP_scale/Aimless for two crystals (**Table 2**). The final stages of refinement were challenged by the disordered regions of the N-terminal domains with final model B-factors at the high end of the distribution when compared to others structures of similar resolution. This is likely due to ∼25% of the molecular mass within the crystal unable to be included in the final model. While the majority of the refinement stages were performed within Phenix using the spherically truncated data from Proteum3, issues during deposition validation prompted a systematic comparison of data processing and final refinement programs, part of which is shown in **Table 2**. The first four columns in the table show the final iteration of refinement for the same input molecule by either Phenix or Refmac utilizing data processed with Proteum3 or aP_scale/Aimless. Upon deposition validation, significant variant, most notably in RSRZ, were observed. The final model refined has been deposited with PDB ID 6VJI and includes the associated spherically truncated structure factors. The deposit also includes the native crystal data, selenomethionine peak wavelength, iodide derivative and gold derivative ellipsoidally truncated used for phasing along with data collected at the zinc peak wavelength. Raw diffraction data will also be made available.

**Table 2.**
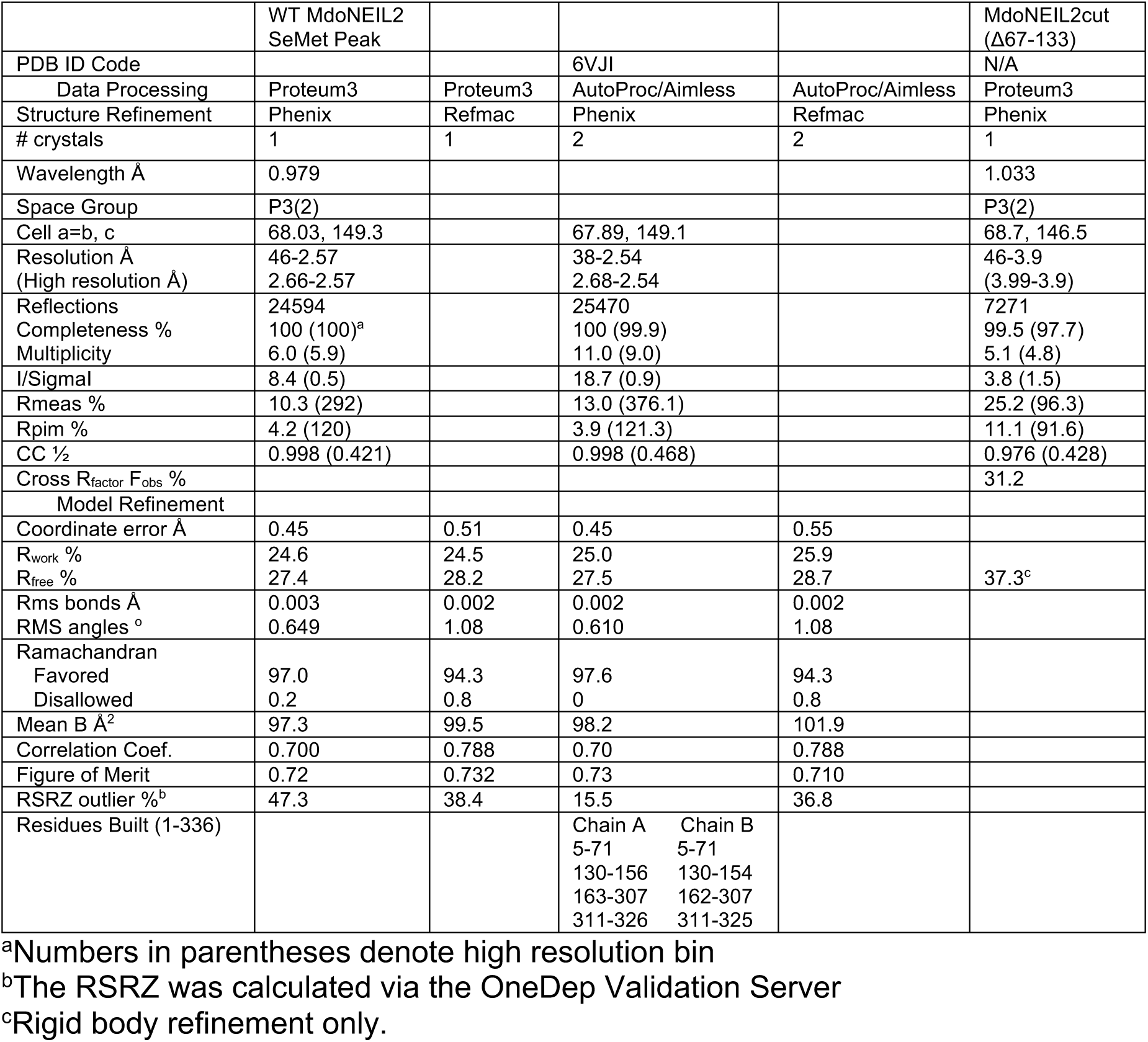
Crystallographic data processing using spherically truncated data and model statistics for structure refinement of MdoNEIL2 using multiple programs. Collection statistics are also shown for the construct where the large disordered region of the N-terminus was removed. The deposited coordinates are from the column with the PDB ID code.

The truncated form of MdoNEIL2 was evaluated via both isomorphous differences (Phenix) and molecular replacement (Phaser(21)), applying search routines with the N and C terminal domains separately. Due to the low resolution, only rigid body refinement was evaluated using Phenix (**Table 2**).

### SAXS data collection and interpretation

Samples for SAXS analysis performed in batch/static mode were prepared by additional purification over a Superdex S200 gel filtration column (GE Life Sciences Inc.) in 25 mM HEPES pH 7.5, 100 mM NaCl, 1 mM DTT and 2-4% glycerol with the equivalent buffer utilized for background subtraction. A concentration series was screened for each protein to evaluate concentration-dependent oligomerization. Data were collected at the SIBYLS beamline (Advanced Light Source, Lawrence Berkeley National Laboratory) through the mail-in data collection program at 11-12 KeV at 0.3-0.5 sec/frame for a 10-15 second run on a Pilatus 6M detector (**Table 3**). Samples for in-line SEC-SAXS were run in 25 mM BisTris pH 8, 150 mM NaCl, 2% glycerol and 1 mM TCEP using a Shodex PROTEIN KW-802.5 at 12 KeV and 3 sec/frame. Background regions were selected and corrections were performed using Chromixs(22). All SAXS primary data analysis was performed using ATSAS 2.8.4(23) with Guinier approximation performed with PRIMUS(24). Fitting of scattering curves to theoretical scattering based on deposited crystal structures was performed using FoXS(25). The EcoNei protein used for SAXS experiments was expressed and purified as described in(26) while the expression and purification protocol of human PCNA (used as control) was previously described in(27). The data utilized for SAXS analysis will be made available via the Small Angle Scattering Biological Data Bank (SASDB) (*www.sasbdb.org).*

**Table 3.** SAXS data collection and analysis parameters.

| **Data collection parameters** | PCNA | EcoNei | EcoNei SEC | MdoNEIL2 | MdoNEIL2 SEC | MdoNEIL2cut SEC |
| --- | --- | --- | --- | --- | --- | --- |
|  | 160323 | 160524 | 180309 | 160323 | 180309 | 180309 |
| Wavelength (Å) | 1.127 | 1.127 | 1.03 | 1.127 | 1.03 | 1.03 |
| Q Range (Å<sup>-1</sup>) | 0.013 – 0.512 | 0.017 – 0.323 | 0.016 – 0.344 | 0.019 – 0.320 | 0.015 – 0.323 | 0.016 – 0.371 |
| Exposure time (s) | 0.5 sec/15 | 0.3 sec/10 | 3 sec | 0.5 sec/15 | 3 sec | 3 sec |
| Concentration (mg/ml) | 3.2 – 8.5 | 0.7 – 2.3 | 5 | 2.6-4.2 | 5 | 10 |
| **Structural parameters** |  |  |  |  |  |  |
| $I(0)$ from Guinier | 1823 ± 4.8 | 113.1 ± 1.7 | 264.9 ± 0.7 | 489.9 ± 3.6 | 185.6 ± 0.9 | 353.1 ± 0.9 |
| $R_g$ (Å) from Guinier | 34.6 ± 0.1 | 24.2 ± 0.5 | 23.3 ± 0.9 | 30.9 ± 0.3 | 26.6 ± 0.6 | 24.4 ± 0.1 |
| $I(0)$ from $P(r)$ | 1817 | 114.4 | 268.0 | 493.8 | 185.2 | 354.8 |
| $R_g$ (Å) from $P(r)$ | 34.2 | 25.0 | 24.0 | 32.1 | 26.9 | 24.9 |
| $D_{max}$ (Å) | 105 | 81.9 | 78.0 | 112.2 | 82.0 | 78.3 |
| Porod volume (Å<sup>3</sup>) | 132000 | 48300 | 44600 | 77600 | 57800 | 46800 |
| **Molecular weight determination (kDa)** |  |  |  |  |  |  |
| MW<sub>SAXSMoW</sub> | 88.3 | 28.6 | 30.4 | 57.1 | 41.1 | 30.6 |
| Expected theoretical | 89.1 | 30.7 | 30.7 | 39.0 | 39.0 | 32.7 |

## RESULTS

### Crystallization and Structure Determination

Human NEIL2 (HsaNEIL2) contains 332 residues with sequence analysis predicting large disorder content between residues 55 (immediately after β-strand 2) and 125. Not surprisingly full-length HsaNEIL2 failed to crystallize. Additionally, HsaNEIL2 was problematic for solution studies due to multimodal polydispersity in dynamic light scattering experiments; aggregation and radiation damage were also observed in SAXS experiments. The N-terminal domain of the Fpg/Nei structural family contains 8 β-strands and secondary structure predictions using HsaNEIL2 and several orthologs were inconclusive with regards to the boundaries for proper truncation of the disordered region due to predicted β-strand segments within that region for some orthologs. Two orthologs with predicted lower disorder content in the N-terminal domain were cloned, expressed, purified and crystallized: The ortholog from *Xenopus tropicalis* produced crystalline needle clusters that failed to optimize whereas the mammalian ortholog from gray short tailed opossum *Monodelphis domestica* yielded diffraction-quality single crystals.

With the low sequence conservation within the Fpg/Nei structural family and the symmetrical nature of N-terminal fold resulting in a 4-fold ambiguity in secondary structure matching, structure determination using molecular replacement was unsuccessful and necessitated experimental phasing methods. Numerous trials for SAD and MAD utilizing selenomethionine-substituted protein as well as SIRAS methods were explored and produced partial phase solutions that stalled during building and refinement. The combination of Se-MAD and MIRAS as well as phasing using anisotropically processed data verified a strong non-crystallographic symmetry axis within Laue group 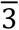 parallel to that of a crystallographic axis for the higher ordered Laue group 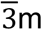 (**Supplemental Figure 1)**. The orientation of the axis mimicked the P3_2_12/P3_1_12 spaces groups with a combined ambiguity of the screw axis. The SAD/MIR phases resolved the final space group to P3_2_ with 2 molecules in the asymmetric unit. The solution included native data, Se-SAD at peak wavelength, along with iodide and gold derivatives. A native crystal collected at the peak wavelength for Zn was utilized for verification of the ion in the zinc finger motif, which displays anomalous signal at all wavelengths used for phasing.

Mutation of L7, 210 and 284 to Met with subsequent data collection of the selenomethionine derivative at peak wavelength was used to verify their locations. They were chosen strategically as L7 is in the N-terminal helix, L210 lies in the interdomain linker and L284 is in the C-terminal domain away from the non-crystallographic two-fold. Final data and model analysis revealed the hurdles that had plagued the phase determination using the Se-MAD data: Of the 4 endogenous Met residues in the wild-type protein, M39 displayed broad and diffuse anomalous signal indicating two conformations, M123 is located in a disordered region, and M301 lies directly across the NCS two-fold rotation axis.

### Overall structure of NEIL2

The structure of the MdoNEIL2 was determined to 2.54 Å with R_free_ 27.5 % and crystallized with two molecules in the asymmetric unit related by a non-crystallographic two-fold axis. The final model for the two chains contains 251 and 255 amino acids of the 336 amino acid sequence with the primary omitted section being the large internal NEIL2 insert (residues 66-131). Some residual density was observed for this region but its position along the non-crystallographic two-fold negated its inclusion due to ambiguity (**Supplemental Figure 1**). A pair of weakly anomalous scattering peaks could be observed for both the WT and L7/210/284M selenomethionine data, indicating partial ordering of M123 in the insert region. The two chains in the asymmetric unit show a 0.25 Å RMS for the C-alpha trace with only minor differences between the two chains in N-terminal domain loops and the respective refined B-factors (**Supplemental Figure 1**). For simplicity, the remainder of the description of the structure will be limited to a single chain (Chain A).

The general fold of NEIL2 is composed of an N-terminal β-sandwich domain and a C-terminal domain containing the H2TH and Zinc finger DNA-binding motifs typical of the Fpg/Nei superfamily of glycosylases (**Figure 2**)(28). Unique to NEIL2 is an extended interdomain linker along with a large insert within the N-terminal domain (aa 66-131 in opNei2). The positions of methionines 36, 39 and 301 in the WT protein were identified using anomalous scattering from the selenomethionine-substituted protein crystals as were the site-directed mutagenesis of leucines at positions 7, 210 and 284 (13). The heavy atom soaks provided identification of cysteines 183 and 284 (**Figure 2A,B**) while the zinc atom in the predicted CHCC-type zinc finger was verified by X-ray fluorescence at the peak wavelength for zinc.

The core portions of the N- and C-terminal domains were compared to other proteins in the Fpg/Nei, with the N-terminal domain showing a 1.75 Å RMS deviation (22% sequence identity over 92 residues, 377 atoms) and 0.78 Å RMSD (29% sequence identity over 100 residues, 483 atoms) for the N- and C-terminal domains of mouse NEIL3, respectively. The N-terminal domain shares very little sequence conservation with other glycosylases of the same family outside of the N-terminal helix active site P2-E3-G4 motif and conserved lysine (K50 in MdoNeil2 and HsaNEIL2), thus hindering structure solution via molecular replacement. As the large insert in NEIL2 contains significant disorder preventing its inclusion in the final model and resulting elevated B-factors, a protein construct was engineered with the region omitted. Omission of this region had little impact on enzymatic activity (below), however the construct proved more difficult to crystallize with the diffraction limit reaching only to 4.3 Å, with pronounced anisotropic diffraction. The crystals were of the same space group with minimal change in unit cell parameters. Isomorphous difference methods with a R_merge_ of 31% for F_obs_ between full-length and truncated data sets (**Table 1, Supplemental Figure 2**), a 34.5% R_free_ with the isomorphous replacement model upon rigid body refinement, and the molecular replacement solution all show the truncated protein to be in the same conformation as the full-length protein, suggesting that the insert aided in crystal growth but does not influence the overall structure of the protein within the crystal.

### The NEIL2 ortholog is active on both single and double-stranded DNA

The activity of NEIL2 was assessed using cyanoborohydride trapping and glycosylase activity assays (**Figure 3, Supplemental Figure 3)** The opossum ortholog shows 57% identity and 70% homology with the human enzyme with the variability in sequence of the unique insert accounting for 10% of the divergence. Not surprisingly, the human and opossum ortholog showed a similar substrate profile using the trapping assay with greater activity towards single-stranded DNA substrates. NEIL2 shows modest activity towards lesions within duplex DNA. However, unlike what was observed for NEIL3, NEIL2 shows robust activity for the apurinic/apyrimidinic (AP) site within duplex DNA. In both single and double-stranded contexts, glycosylase activity reaches completion with enzyme to substrate ratios in the 4:1 to 16:1 range for both the full-length protein and the construct with the disordered region removed. The abasic site was the preferred substrate in both contexts. For the oxidized base lesions, dHU showed the greatest activity within single-stranded DNA reaching completion at the highest enzyme ratio while OHU showed the greatest activity in double-stranded DNA but only reached ∼30% completion under the conditions employed. The poor Tg activity along with AP site preference are in agreement with what was previously reported (29). We observed in our assays that the primary product of the reaction for single-stranded DNA appears to be the δ-elimination product with the β-elimination product being favored for duplex DNA substrates (**Figure 3, Supplemental Figure 3**).

### Comparison with other Fpg/Nei glycosylases

The NEIL2 structure revealed, unexpectedly, that the orientation of the C-terminal domain relative to the N-terminal domain differs significantly from that of other proteins in the family (**Figure 2C**). While NEIL1 adopts a “closed” conformation whether it is bound to DNA or not, NEIL2 exhibits an “open” conformation, which places the N-terminal catalytic residue on the opposite face of the C-terminal domain DNA-binding motifs requiring a near 80° rotation (**Figure 4**) of the domains relative to each other to achieve catalytic competency. The only other time this has been observed for proteins in this family is for the more distantly related endonuclease VIII glycosylase from *E. coli* (EcoNei) (**Figure 4A**)(30, 31). The NEIL glycosylases differ from their bacterial counterparts because they contain long disordered regions(32). While NEIL1 and NEIL3 harbor these flexible extensions at their C-terminus sequence alignments revealed that NEIL2 is unique in that the disordered region is located within the glycosylase fold, more precisely in the N-terminal domain. Two interesting unique NEIL2 features are revealed within the N-terminal domain pertaining to insertion elements: a largely disordered region (aa 65-130) resides between β strands 3 and 4 and, in the “open” conformation projects towards the typical DNA binding interface with the C-terminal domain. A second, smaller insert of 8-10 residues, also unique to NEIL2, is located between β strands 5 and 6 and contains 3 lysines and 1 arginine in the opossum ortholog, and 3 lysines and 2 arginines in the human sequence. The N-terminal domain rotation expected for assembly of a catalytically competent complex would put this smaller insert in proximity to the DNA substrate binding cleft (**Figure 5**).

**Figure 4.**
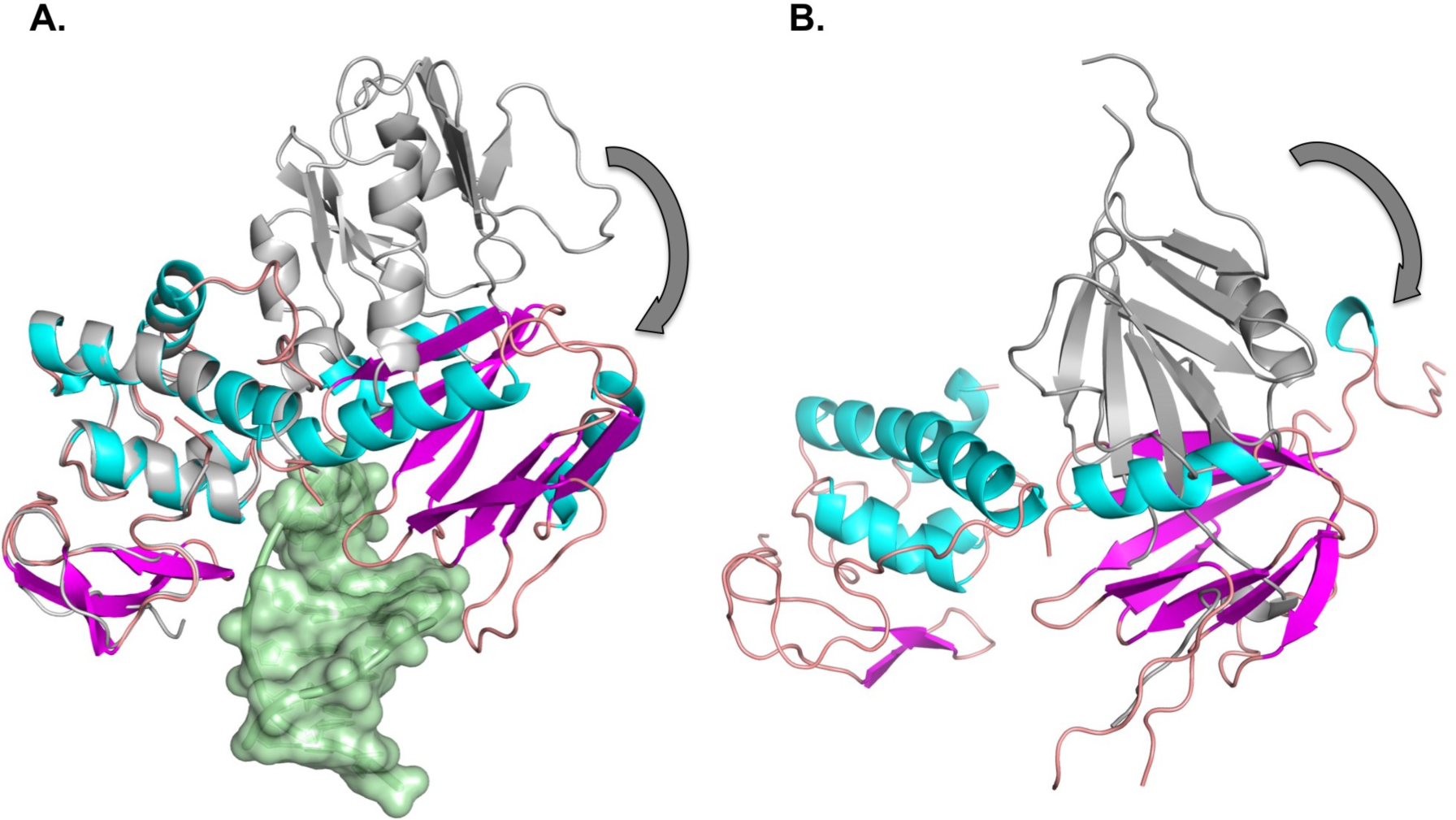
Comparison of the open and closed forms of *E. coli* endonuclease VIII (EcoNei) with MdoNEIL2. A.) The unliganded (grey; PDB ID code: 1Q3B)(31) and DNA-bound (shown in secondary structure colors: helices in cyan and β strands in magenta; PDB ID code 2EA0) (Golan, Zharov, Grollman & Shoham, unpublished 2007) forms of EcoNei were overlaid based on their C-terminal domains. A significant rotation (reported to be ∼50°) of the N-terminal domain for the unliganded structure would be required for catalytic competency(31). B.) The MdoNEIL2 structure (grey) is overlaid with the expected conformation of a catalytically competent complex (secondary structure colors). An angle of ∼80° for rotation of the NEIL2 N-terminal domain between the two conformations was determined using Superpose within CCP4(48). The interdomain linker for MdoNEIL2 has been omitted for clarity.

**Figure 5.**
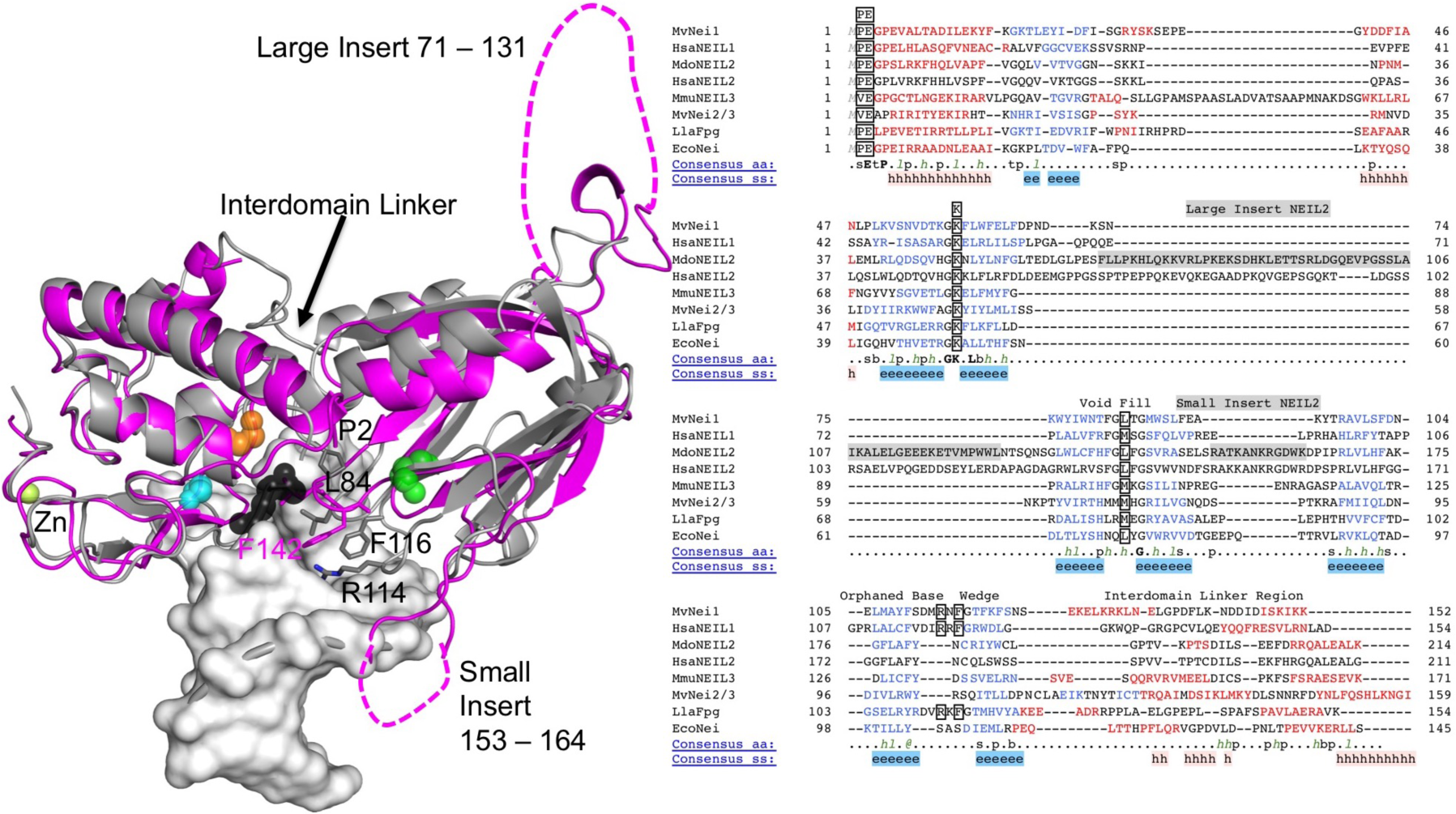
Comparison using least-squares superposition of the N-terminal domain of proteins in the Fpg/Nei superfamily. Crystal structure of the mimivirus NEIL1 ortholog (33) in complex with abasic site containing duplex, colored grey, with key loops and the respective residues involved in catalytic function shown: (P2, E3), void-filling residues (L84, R114 and F116). The black spheres represent the abasic site lesion. The MdoNEIL2 structure is shown in magenta upon independent superposition of the N-terminal and C-terminal domains. Residues corresponding to HsaNEIL2 cancer variants S140 (green), G230 (orange), and G303 (cyan) are highlighted. The structure-based sequence alignment for the N-terminal domain of the Fpg/Nei family was produced using PROMALS3D(49). While the loop between β strands 4 and 5 containing the void-filling residue is consistent in length within the Fpg/Nei family, the loop between β strands 7 and 8 is quite variable, with the NEIL2/3 orthologs being significantly shorter. The structures used in the alignment are MvNei1 (PDB ID code: 3A46)(33), HsaNEIL1 (1TDH)(3), MmuNEIL3 (3W0F)(34), MvNei2/3 (4MB7)(50), LlaFpg (1L1T)(51), and EcoNei (2EA0). The full structure-based sequence alignment is provided in **Supplemental Figure 4**.

Despite an overall low sequence conservation NEIL2 retains some key features within the N-terminal domain compared to other glycosylases of the Fpg/Nei superfamily: an N-terminal active site (**Figure 5, Supplemental Figure 4**), a conserved lysine critical to activity located in a short loop between β strands 2 and 3 (Lys50 in MdoNeil2/HsaNEIL2 vs. Lys54 in HsaNEIL1), and a loop between β strands 4 and 5 containing a hydrophobic residue (typically Leu or Met) that stabilizes the everted damaged base into the active site of the enzyme(33). Because of the large insert in NEIL2, prediction of the location for the β4-β5 loop was uncertain. The structure of MdoNEIL2 determined here reveals that the β4-β5 loop comprises residues 140-143, with Leu141 and Phe142 as candidates to serve the function of stabilizing the damaged base everted into the active site. The position of a third loop, located between β strands 7 and 8, which in NEIL1 harbors two residues stabilizing the orphaned base on the non-damaged strand (typically an Arg and a Tyr/Phe) was equally uncertain. Similar to what was observed for the ssDNA-dependent NEIL3, the β7-β8 loop containing two of the void-filling residues in NEIL1 is significantly shorter for NEIL2(34), NEIL2 therefore lacks two of the void-filling residues, implying that lesion search and recognition will differ from what was described for NEIL1(33, 35, 36).

### Conformational dynamics in solution

The observation of the unique interdomain conformation of NEIL2 prompted further investigation into the solution behavior of the protein utilizing SAXS (**Table 3, SAXS 6**). While crystal structures exist for *E. coli* endonuclease VIII (EcoNei) showing distinct conformations of the enzyme between unliganded and bound to DNA(30, 31), this behavior has not been investigated in solution. We conducted SAXS experiments for the unliganded protein to see if the structure observed in the crystal represents the solution state of the enzyme (**Figure 6B**). Data were collected under static conditions as well as utilizing in-line size exclusion chromatography with the subsequent scattering curves fit to the theoretical curves generated from the published crystal structures, using FoXS and utilizing PCNA as a method control (**Figure 6A**). Both methods provide strong evidence that the conformation observed for the unliganded EcoNei represents the state observed in solution, suggesting that this enzyme undergoes a specific conformational event upon binding DNA. Evaluation of particle shape in the form of Rg and Dmax shows PCNA to fall within values predicted from the crystal structures as well as in agreement with previous SAXS studies(27). While no SAXS studies have been to date published for EcoNei, vales for Rg and Dmax were consistent with the crystal structure.

**Figure 6.**
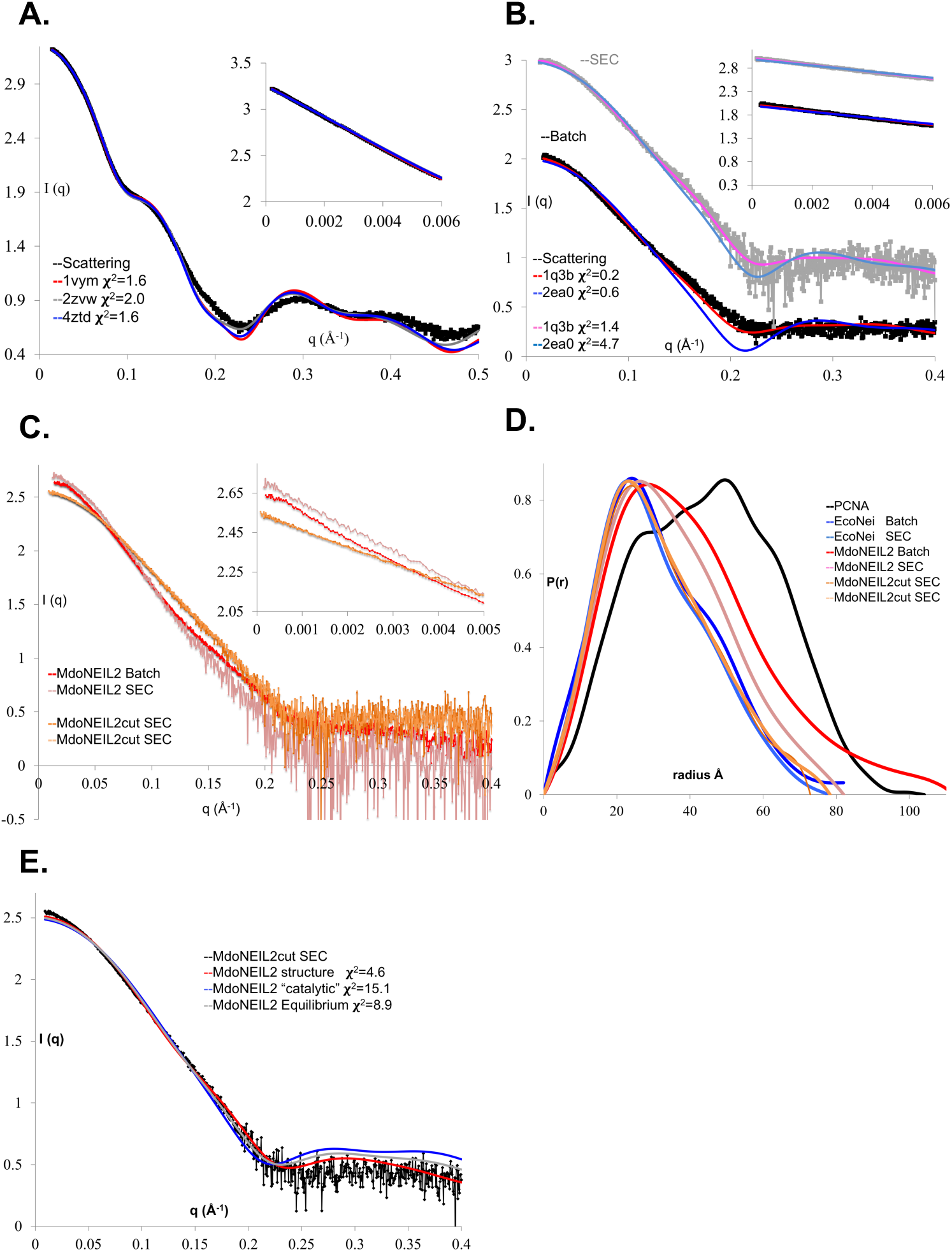
SAXS analysis for PCNA, EcoNei, MdoNEIL2 and MdoNEIL2cut. A.) Shown is the scattering curve for PCNA (experimental control) and fits to 3 crystal structures, after omission of any ligands, using FoXS(25). B.) Shown is the SAXS profile for EcoNei under batch conditions (black) or upon elution from in-line size exclusion chromatography (grey). The fits shown represent the apo conformation (red and pink) and DNA bound (dark and light blue). C.) Scattering curves for batch and SEC-SAXS for full-length MdoNEIL2 (red and pink) and SEC-SAXS for MdoNEIL2cut (orange). For panels A, B and C, the inset shows the Guinier region. D.) Distance distribution function for all samples. E.) FoXS fitting for the MdoNEIL2cut.

Both full-length and insert-deleted NEIL2 (MdoNEIL2cut) were evaluated, with the latter only amenable to the in-line SEC data collection due to instability within batch methods (**Figure 6C**). Size exclusion chromatography, dynamic light scattering, and multi-angle light scattering showed both constructs to be homogeneous monomeric populations in solution. The full-length NEIL2 showed Rg as well as pair-wise distance distribution functions consistent with being larger than EcoNei. However, the Dmax, particularly under batch conditions, was considerably larger (**Figure 6D**). Like the MdoNEIL2cut construct, the full-length protein showed improvement by application of in-line SEC methods. The distance distribution profile, like the primary scattering curve for MdoNEIL2cut was quite similar to that observed for EcoNei indicating some agreement in particle shape. As the large disordered insert cannot directly be fit to the scattering curve of the full-length protein, structure fitting was only performed using the MdoNEIL2cut data. Like that of EcoNei, the best fit to data could be achieved to the crystal structure and not to a model produced by allowing the N- and C-terminal domains to rotate to a catalytically competent conformation (**Figure 6E**). However, the quality of the fit in the low Q and Porod regions deviates significantly from the experimental curve compared to that of the PCNA control or EcoNei. This finding indicates that neither the MdoNEIL2 apo structure, the expected catalytic conformation, nor an average between the two theoretical curves (representing equilibrium of the two conformations) represents the true behavior of MdoNEIL2 in solution.

The data for both EcoNei and MdoNEIL2 were further evaluated with MultiFoXS, an ensembling method not restricted to the input conformation of the protein. Both proteins were allowed flexibility in the linker (residues 125-134 for EcoNei and 190-214 for MdoNEIL2) and evaluated for up to 5 states with the final selection of 10 models for each state (**Supplemental Figure 5**). The method suggested between 2 and 3 states to provide the best fit to the data. For EcoNei, the input unliganded structure was selected for 3 of the 10 models for conformation 1 and in none of the models produced for either of the 5 states was the catalytically competent conformation produced. Conversely, neither the crystal structure conformation nor the conformation predicted for catalytic competency were sampled during the simulation. Additional analysis of the particle envelopes using Dammif displayed convergence to a discrete population for EcoNei but highly variable for MdoNEIL2 (**Supplemental Figure 5**). Overall the data suggest that while EcoNei likely exists in discrete populations, MdoNEIL2 is free to sample more conformational space in the absence of DNA.

### Analysis of HsaNEIL2 cancer variants

Several variants were selected from The Cancer Genome Atlas (TCGA) by the UVM Bioinformatics Shared Resource for evaluation of biochemical function guided by the crystal structure of MdoNEIL2. G230W, G233 in MdoNEIL2, showed a significant defect in activity (**Figure 3B**), predominantly with ssDNA. This glycine is the N-terminal cap of a helix in the H2TH motif of the C-terminal domain. This residue is absolutely conserved (**Supplemental Figure 4**) within the Fpg/Nei family. It provides the N-terminal cap of the helix coordinating the catalytic residue Glu3, and the adjacent residue, Asn231, is also strictly conserved as it forms a hydrogen bond with a non-bridging oxygen of the phosphate backbone 5’ to the lesion. Introducing a large aromatic residue like tryptophan at the location of G230 is predicted to induce local changes that would negatively affect DNA binding and activity. A second variant, G303R in the zinc finger motif, also showed a significant decrease in glycosylase activity, albeit less severe compared with G230W, whereas lyase activity was largely retained for both ssDNA and dsDNA. Interestingly, the assay product observed for ssDNA for the variants was β-elimination, like that seen for dsDNA substrate and not δ-elimination (**Supplemental Figure 3**). A third variant, S140N was also evaluated. It resides at the N-terminal end of β-strand 5, which resides between the canonical void-filling loop and the small insert in NEIL2. S140N showed no change in overall glycosylase activity relative to wild-type but appeared to follow the trend of the other variants, with a modest shift in the equilibrium between β-elimination and δ-elimination (**Supplemental Figure 3E**).

## DISCUSSION

Mammalian DNA is a dynamic and crowded space with numerous processes occurring both sequentially and simultaneously. Coordination of DNA replication, sensing of damage sites for repair and changes in methylation patterns for transcription of activated genes requires the orchestration of recruitment for multifunctional protein factors for use in specific functions. Increasing evidence has placed NEIL2 at the junction of several of these processes through identification of binding partners, their influence on activity and the range of substrates suitable for activity. Its fundamental activity of recognition and removal of oxidized bases and abasic sites within unique structures of DNA is temporally and spatially dependent on the specific function. Currently identified binding partners of NEIL2 include BER and small patch repair DNA polymerase β and XRCC1(1, 37), transcription-associated repair CSB(8), APE1, TDG(11) and represent different steps in DNA repair and maintenance. Here we present the first crystal structure of a NEIL2 glycosylase. The protein adopts an unprecedented “open” conformation not observed in NEIL1 and NEIL3. In that open conformation the large insert in NEIL2 (residues 66-131) juts out of the protein away from the DNA binding interface. We demonstrate that NEIL2 is conformationally dynamic and suggest that the flexibility of NEIL2 allows tethering to other proteins without impeding its interactions with the DNA. This allows for utilization of NEIL2 in alternate environments. Placement of the protein-interaction module within the N-terminal domain of NEIL2 indicates that the repair environment or dynamic interplay between BER partners differs from that of NEIL1, which contains a long unstructured extension to serve a similar function(38), but at its C-terminus.

The crystal structure also revealed that NEIL2 lacks two of the three void-filling or intercalating residues (Arg and Tyr) observed in other glycosylases of the same family (Leu/Met, Arg, Phe/Tyr) (*e.g.*, Met-81 on the β4-β5 loop; Arg-118 and Phe-120 on the β7-β8 loop in human NEIL1)(3, 34, 36).The Leu/Met hydrophobic residue stabilizes the flipped out lesion everted from the double helix. The Arg and Phe/Tyr residues invade the DNA double helix on the side of the orphaned base. The aromatic residue in the triad has been referred to as the wedge residue, because of its role in lesion search. An *E. coli* Fpg variant where Phe-111 was mutated to alanine displayed a reduced glycosylase activity on lesion-containing ds DNA compared to the wild-type enzyme. This decrease in activity could be attributed to a reduced ability to search for lesions(39). The Arg and Phe/Tyr residues are absent in NEIL2 because the loop they reside on is much shorter than that in NEIL1. We described the same situation in the NEIL3 structure(34). Because NEIL2 and NEIL3 prefer ssDNA over ds DNA it is not all that surprising that they lack the loop that harbor residues whose function is to buttress the orphaned base on the opposite strand. But this leaves open the question as to how NEIL2 and NEIL3 search for lesions and which residue, if any, serves as the “wedge” residue.

Of the currently determined structures of enzymes within the Fpg/Nei family, NEIL2 represents only the second example of a glycosylase undergoing large-scale conformational changes. Numerous studies have investigated the conformational events in glycosylases as observed by changes in tryptophan fluorescence including for NEIL1, which experiences small scale dynamics(40) and EcoNei(41) which shows large scale dynamics. These studies can only measure solvent environment change about the Trp residues and cannot speak of distance in the manner that FRET can but both provide time-scale comparisons. Interestingly, the rate constant of the initial measurable step for both enzymes is extremely fast and on equivalent scales at 180×10^6^ s^-1^ for NEIL1 and 12×10^6^ s^-1^ for EcoNei when comparing abasic site substrates. These results follow the structural distinctions that the enzyme with the smallest conformational change (NEIL1) having the faster rate for this step by about 10-fold. However, this must be taken into context by considering the overall rate of the reaction (chemistry) is only 0.1-0.5 s^-1^. In fact, the additional observed microscopic rate constants were quite similar for the two proteins. These results suggest that the initial binding and scanning of the DNA occurs seven orders of magnitude faster than the actual rate of chemistry, regardless of the conformational distance traveled. It is important in the case of NEIL2, which we predict must experience a large domain rearrangement to assemble the catalytically-competent complex in the manner observed for EcoNei, that the conformational step not constitute a functional or kinetic barrier.

Mouse knockdown models of NEIL1,2 resulting in metabolic stress and inflammation but modest or no elevation in mutational load(5) suggest a substrate repertoire that overlaps with other enzymes or that proteins like NEIL2 serve functions outside of basal genome maintenance. The strong AP site activity on both single and double-stranded DNA suggests that either the abasic site is a true substrate for NEIL2, the favored base modification substrate is as of yet unidentified, or sequence-context dependence for a particular lesion may exist. Indeed, it was recently shown that both EcoNei and Fpg will excise N4,5-dimethylcytosine while leaving 5-methylcytosine and 4-methylcytosine uncut(42). Additionally, both NEIL1 and NEIL2 were shown to function in TDG-mediated turnover of 5hmC(11) at sites of TET dependent DNA demethylation while 5hmC accumulated at sites of DNA damage (43) and thus exhibit the ability to substitute for APE1 in the demethylation pathway. Interestingly, knock down of NEILs resulted in elevated 5fC and 5caC levels whereas APE1 knockdown did not. The proposed association of NEIL2 with transcription-associated repair (6), juxtaposed with the difficulty in producing cancer phenotypes in mice(5), suggests that the true function of NEIL2 may yet to be established. Transient sources of bubble or transient ssDNA occur during transcription and recombination events. While errors in replicated DNA strands may be tracked experimentally, sites left unrepaired during translation resulting in aberrant protein synthesis is more difficult to measure but contribute to inflammation.

Recent studies implicate a potential role for NEIL2 in cytosine deaminase APOBEC3-mediated mutagenesis, likely via its activity towards abasic sites(44) where data suggest NEIL2 may outcompete APE1 for the site in particular contexts. This presents interesting contribution of NEIL2 variants to the APOBEC3 mutation signature both by aberrant NEIL2 activity towards the abasic sites produced during the deamination process as well as reduced activity towards oxidized cytosine lesions. The variants studied here displayed the most significant decrease in activity for the cytosine derived substrates 5-OHU in both single- and double-stranded DNA contexts with DHU in the single-stranded substrate. The variant G230W also showed a pronounced decrease in activity for the abasic site in both contexts. Recent data also suggest that both NEIL1 and NEIL2 show preference for a D-loop mimic over the single-stranded exposed lesion within bubble structure substrates(45) while displaying differential activity depending on the location of the lesion within the bubble. Moreover, removal of hydantoin lesions by NEIL1 was shown to be context-dependent(46). The structural and biochemical investigations presented here provide insight into the dynamic behavior of NEIL2. Our work also illuminates how variants of the enzyme can contribute to disease through either mutagenesis-based genomic instability or by influencing gene expression through action on unique DNA structures encountered during transcription.

## Supporting information

Supplemental Infomation

## ACKNOWLEDGEMENTS

Single crystal X-ray diffraction experiments were conducted at GM/CA@APS which has been funded in whole or in part with Federal funds from the National Cancer Institute (ACB-12002) and the National Institute of General Medical Sciences (AGM-12006). This research used resources of the Advanced Photon Source, a U.S. Department of Energy (DOE) Office of Science User Facility operated for the DOE Office of Science by Argonne National Laboratory under Contract No. DE-AC02-06CH11357. The SAXS data collections were conducted at the Advanced Light Source (ALS), a national user facility operated by Lawrence Berkeley National Laboratory on behalf of the Department of Energy, Office of Basic Energy Sciences, through the Integrated Diffraction Analysis Technologies (IDAT) program, supported by DOE Office of Biological and Environmental Research. Additional support comes from the National Institute of Health project MINOS (R01GM105404) and a High-End Instrumentation Grant S10OD018483. SAXS data was collected at SIBYLS which is funded by DOE/BER grant Integrated Diffraction Analysis (IDAT) grant contract number DE-AC02-05CH11231 and NIH MINOS RO1. Identification and selection of human variants along with DNA sequencing services were performed using the Bioinformatics Shared Resource and Vermont Integrative Genomics Resource at the University of Vermont. The authors thank Clemens Vonrhein of Global Phasing Limited for assistance with STARANISO data processing and SHARP/AutoSharp.

## AUTHOR CONTRIBUTIONS

B.E.E. and S.D. wrote the manuscript. S.D. designed the project. B.E.E. designed and performed experiments including protein expression and purification, crystallizations, processing of crystal and solution diffraction data, solution and refinement of structures, activity assays. A.M.A. expressed and purified proteins. V.C. expressed and purified proteins and helped with the activity assays. J.A.D. performed analysis of TCGA for variant selection.

## DECLARATION OF INTERESTS

The authors declare no competing interests.

## Notes

Funding sources: This work was supported by NIH grant P01 CA098993 awarded to S.D. B.E.E. is supported by NIH grant R50 CA233185.

## REFERENCES

1. Das A, Wiederhold L, Leppard JB, Kedar P, Prasad R, Wang H, Boldogh I, Karimi-Busheri F, Weinfeld M, Tomkinson AE, Wilson SH, Mitra S, Hazra TK. NEIL2-initiated, APE-independent repair of oxidized bases in DNA: Evidence for a repair complex in human cells. DNA Repair (Amst). 2006;5(12):1439-48. Epub 2006/09/20. doi: 10.1016/j.dnarep.2006.07.003. PubMed PMID: 16982218; PMCID: PMC2805168.

2. Hazra TK, Kow YW, Hatahet Z, Imhoff B, Boldogh I, Mokkapati SK, Mitra S, Izumi T. Identification and characterization of a novel human DNA glycosylase for repair of cytosine-derived lesions. J Biol Chem. 2002;277(34):30417-20. Epub 2002/07/05. doi: 10.1074/jbc.C200355200. PubMed PMID: 12097317.

3. Doublié S, Bandaru V, Bond JP, Wallace SS. The crystal structure of human endonuclease VIII-like 1 (NEIL1) reveals a zincless finger motif required for glycosylase activity. Proc Natl Acad Sci U S A. 2004;101(28):10284-9. Epub 2004/07/03. doi: 10.1073/pnas.0402051101. PubMed PMID: 15232006; PMCID: PMC478564.

4. Mullins EA, Rodriguez AA, Bradley NP, Eichman BF. Emerging Roles of DNA Glycosylases and the Base Excision Repair Pathway. Trends Biochem Sci. 2019;44(9):765-81. Epub 2019/05/13. doi: 10.1016/j.tibs.2019.04.006. PubMed PMID: 31078398; PMCID: PMC6699911.

5. Rolseth V, Luna L, Olsen AK, Suganthan R, Scheffler K, Neurauter CG, Esbensen Y, Kusnierczyk A, Hildrestrand GA, Graupner A, Andersen JM, Slupphaug G, Klungland A, Nilsen H, Bjoras M. No cancer predisposition or increased spontaneous mutation frequencies in NEIL DNA glycosylases-deficient mice. Sci Rep. 2017;7(1):4384. Epub 2017/07/01. doi: 10.1038/s41598-017-04472-4. PubMed PMID: 28663564; PMCID: PMC5491499.

6. Chakraborty A, Wakamiya M, Venkova-Canova T, Pandita RK, Aguilera-Aguirre L, Sarker AH, Singh DK, Hosoki K, Wood TG, Sharma G, Cardenas V, Sarkar PS, Sur S, Pandita TK, Boldogh I, Hazra TK. Neil2-null Mice Accumulate Oxidized DNA Bases in the Transcriptionally Active Sequences of the Genome and Are Susceptible to Innate Inflammation. J Biol Chem. 2015;290(41):24636-48. Epub 2015/08/08. doi: 10.1074/jbc.M115.658146. PubMed PMID: 26245904; PMCID: PMC4598976.

7. Banerjee D, Mandal SM, Das A, Hegde ML, Das S, Bhakat KK, Boldogh I, Sarkar PS, Mitra S, Hazra TK. Preferential repair of oxidized base damage in the transcribed genes of mammalian cells. J Biol Chem. 2011;286(8):6006-16. Epub 2010/12/21. doi: 10.1074/jbc.M110.198796. PubMed PMID: 21169365; PMCID: PMC3057786.

8. Aamann MD, Hvitby C, Popuri V, Muftuoglu M, Lemminger L, Skeby CK, Keijzers G, Ahn B, Bjoras M, Bohr VA, Stevnsner T. Cockayne Syndrome group B protein stimulates NEIL2 DNA glycosylase activity. Mech Ageing Dev. 2014;135:1-14. Epub 2014/01/11. doi: 10.1016/j.mad.2013.12.008. PubMed PMID: 24406253; PMCID: PMC3954709.

9. Osorio A, Milne RL, Kuchenbaecker K, Vaclova T, Pita G, Alonso R, Peterlongo P, Blanco I, de la Hoya M, Duran M, Diez O, Ramon YCT, Konstantopoulou I, Martinez-Bouzas C, Andres Conejero R, Soucy P, McGuffog L, Barrowdale D, Lee A, Swe B, Arver B, Rantala J, Loman N, Ehrencrona H, Olopade OI, Beattie MS, Domchek SM, Nathanson K, Rebbeck TR, Arun BK, Karlan BY, Walsh C, Lester J, John EM, Whittemore AS, Daly MB, Southey M, Hopper J, Terry MB, Buys SS, Janavicius R, Dorfling CM, van Rensburg EJ, Steele L, Neuhausen SL, Ding YC, Hansen TV, Jonson L, Ejlertsen B, Gerdes AM, Infante M, Herraez B, Moreno LT, Weitzel JN, Herzog J, Weeman K, Manoukian S, Peissel B, Zaffaroni D, Scuvera G, Bonanni B, Mariette F, Volorio S, Viel A, Varesco L, Papi L, Ottini L, Tibiletti MG, Radice P, Yannoukakos D, Garber J, Ellis S, Frost D, Platte R, Fineberg E, Evans G, Lalloo F, Izatt L, Eeles R, Adlard J, Davidson R, Cole T, Eccles D, Cook J, Hodgson S, Brewer C, Tischkowitz M, Douglas F, Porteous M, Side L, Walker L, Morrison P, Donaldson A, Kennedy J, Foo C, Godwin AK, Schmutzler RK, Wappenschmidt B, Rhiem K, Engel C, Meindl A, Ditsch N, Arnold N, Plendl HJ, Niederacher D, Sutter C, Wang-Gohrke S, Steinemann D, Preisler-Adams S, Kast K, Varon-Mateeva R, Gehrig A, Stoppa-Lyonnet D, Sinilnikova OM, Mazoyer S, Damiola F, Poppe B, Claes K, Piedmonte M, Tucker K, Backes F, Rodriguez G, Brewster W, Wakeley K, Rutherford T, Caldes T, Nevanlinna H, Aittomaki K, Rookus MA, van Os TA, van der Kolk L, de Lange JL, Meijers-Heijboer HE, van der Hout AH, van Asperen CJ, Gomez Garcia EB, Hoogerbrugge N, Collee JM, van Deurzen CH, van der Luijt RB, Devilee P, Hebon, Olah E, Lazaro C, Teule A, Menendez M, Jakubowska A, Cybulski C, Gronwald J, Lubinski J, Durda K, Jaworska-Bieniek K, Johannsson OT, Maugard C, Montagna M, Tognazzo S, Teixeira MR, Healey S, Investigators K, Olswold C, Guidugli L, Lindor N, Slager S, Szabo CI, Vijai J, Robson M, Kauff N, Zhang L, Rau-Murthy R, Fink-Retter A, Singer CF, Rappaport C, Geschwantler Kaulich D, Pfeiler G, Tea MK, Berger A, Phelan CM, Greene MH, Mai PL, Lejbkowicz F, Andrulis I, Mulligan AM, Glendon G, Toland AE, Bojesen A, Pedersen IS, Sunde L, Thomassen M, Kruse TA, Jensen UB, Friedman E, Laitman Y, Shimon SP, Simard J, Easton DF, Offit K, Couch FJ, Chenevix-Trench G, Antoniou AC, Benitez J. DNA glycosylases involved in base excision repair may be associated with cancer risk in BRCA1 and BRCA2 mutation carriers. PLoS Genet. 2014;10(4):e1004256. Epub 2014/04/05. doi: 10.1371/journal.pgen.1004256. PubMed PMID: 24698998; PMCID: PMC3974638.

10. Benitez-Buelga C, Baquero JM, Vaclova T, Fernandez V, Martin P, Inglada-Perez L, Urioste M, Osorio A, Benitez J. Genetic variation in the NEIL2 DNA glycosylase gene is associated with oxidative DNA damage in BRCA2 mutation carriers. Oncotarget. 2017;8(70):114626-36. Epub 2018/02/01. doi: 10.18632/oncotarget.22638. PubMed PMID: 29383107; PMCID: PMC5777719.

11. Schomacher L, Han D, Musheev MU, Arab K, Kienhofer S, von Seggern A, Niehrs C. Neil DNA glycosylases promote substrate turnover by Tdg during DNA demethylation. Nat Struct Mol Biol. 2016;23(2):116-24. Epub 2016/01/12. doi: 10.1038/nsmb.3151. PubMed PMID: 26751644; PMCID: PMC4894546.

12. Shayevitch R, Askayo D, Keydar I, Ast G. The importance of DNA methylation of exons on alternative splicing. RNA. 2018;24(10):1351-62. Epub 2018/07/14. doi: 10.1261/rna.064865.117. PubMed PMID: 30002084; PMCID: PMC6140467.

13. Doublié S. Production of selenomethionyl proteins in prokaryotic and eukaryotic expression systems. Methods Mol Biol. 2007;363:91-108. Epub 2007/02/03. doi: 10.1007/978-1-59745-209-0_5. PubMed PMID: 17272838.

14. Prakash A, Carroll BL, Sweasy JB, Wallace SS, Doublié S. Genome and cancer single nucleotide polymorphisms of the human NEIL1 DNA glycosylase: activity, structure, and the effect of editing. DNA Repair (Amst). 2014;14:17-26. Epub 2014/01/03. doi: 10.1016/j.dnarep.2013.12.003. PubMed PMID: 24382305; PMCID: PMC3926126.

15. Kabsch W. Xds. Acta Crystallogr D Biol Crystallogr. 2010;66(Pt 2):125-32. Epub 2010/02/04. doi: 10.1107/S0907444909047337. PubMed PMID: 20124692; PMCID: PMC2815665.

16. Tickle IJ, Flensburg, C., Keller, P., Paciorek, W., Sharff, A., Vonrhein, C., Bricogne, G. STARANISO. Cambridge, United Kingdom: Global Phasing Ltd 2018.

17. Sheldrick GM. A short history of SHELX. Acta Crystallogr A. 2008;64(Pt 1):112-22. Epub 2007/12/25. doi: 10.1107/S0108767307043930. PubMed PMID: 18156677.

18. Vonrhein C, Blanc E, Roversi P, Bricogne G. Automated structure solution with autoSHARP. Methods Mol Biol. 2007;364:215-30. Epub 2006/12/19. doi: 10.1385/1-59745-266-1:215. PubMed PMID: 17172768.

19. Adams PD, Afonine PV, Bunkoczi G, Chen VB, Davis IW, Echols N, Headd JJ, Hung LW, Kapral GJ, Grosse-Kunstleve RW, McCoy AJ, Moriarty NW, Oeffner R, Read RJ, Richardson DC, Richardson JS, Terwilliger TC, Zwart PH. PHENIX: a comprehensive Python-based system for macromolecular structure solution. Acta Crystallogr D Biol Crystallogr. 2010;66(Pt 2):213-21. Epub 2010/02/04. doi: 10.1107/S0907444909052925. PubMed PMID: 20124702; PMCID: PMC2815670.

20. Emsley P, Lohkamp B, Scott WG, Cowtan K. Features and development of Coot. Acta Crystallogr D Biol Crystallogr. 2010;66(Pt 4):486-501. Epub 2010/04/13. doi: 10.1107/S0907444910007493. PubMed PMID: 20383002; PMCID: PMC2852313.

21. McCoy AJ, Grosse-Kunstleve RW, Adams PD, Winn MD, Storoni LC, Read RJ. Phaser crystallographic software. J Appl Crystallogr. 2007;40(Pt 4):658-74. Epub 2007/08/01. doi: 10.1107/S0021889807021206. PubMed PMID: 19461840; PMCID: PMC2483472.

22. Panjkovich A, Svergun DI. CHROMIXS: automatic and interactive analysis of chromatography-coupled small-angle X-ray scattering data. Bioinformatics. 2018;34(11):1944-6. Epub 2018/01/05. doi: 10.1093/bioinformatics/btx846. PubMed PMID: 29300836; PMCID: PMC5972624.

23. Franke D, Petoukhov MV, Konarev PV, Panjkovich A, Tuukkanen A, Mertens HDT, Kikhney AG, Hajizadeh NR, Franklin JM, Jeffries CM, Svergun DI. ATSAS 2.8: a comprehensive data analysis suite for small-angle scattering from macromolecular solutions. J Appl Crystallogr. 2017;50(Pt 4):1212-25. Epub 2017/08/16. doi: 10.1107/S1600576717007786. PubMed PMID: 28808438; PMCID: PMC5541357.

24. Konarev PV, Volkov VV, Sokolova AV, Koch MHJ, Svergun DI. PRIMUS: a Windows PC-based system for small-angle scattering data analysis. Journal of Applied Crystallography. 2003;36:1277–82. doi: 10.1107/S0021889803012779. PubMed PMID: WOS:000185178600026.

25. Schneidman-Duhovny D, Hammel M, Tainer JA, Sali A. FoXS, FoXSDock and MultiFoXS: Single-state and multi-state structural modeling of proteins and their complexes based on SAXS profiles. Nucleic Acids Res. 2016;44(W1):W424-9. Epub 2016/05/07. doi: 10.1093/nar/gkw389. PubMed PMID: 27151198; PMCID: PMC4987932.

26. Jiang D, Hatahet Z, Blaisdell JO, Melamede RJ, Wallace SS. Escherichia coli endonuclease VIII: cloning, sequencing, and overexpression of the nei structural gene and characterization of nei and nei nth mutants. J Bacteriol. 1997;179(11):3773-82. Epub 1997/06/01. PubMed PMID: 9171429; PMCID: PMC179177.

27. Prakash A, Moharana K, Wallace SS, Doublié S. Destabilization of the PCNA trimer mediated by its interaction with the NEIL1 DNA glycosylase. Nucleic Acids Res. 2017;45(5):2897-909. Epub 2016/12/21. doi: 10.1093/nar/gkw1282. PubMed PMID: 27994037; PMCID: PMC5389659.

28. Prakash A, Doublié S, Wallace SS. The Fpg/Nei family of DNA glycosylases: substrates, structures, and search for damage. Prog Mol Biol Transl Sci. 2012;110:71-91. Epub 2012/07/04. doi: 10.1016/B978-0-12-387665-2.00004-3. PubMed PMID: 22749143; PMCID: PMC4101889.

29. Katafuchi A, Nakano T, Masaoka A, Terato H, Iwai S, Hanaoka F, Ide H. Differential specificity of human and Escherichia coli endonuclease III and VIII homologues for oxidative base lesions. J Biol Chem. 2004;279(14):14464-71. Epub 2004/01/22. doi: 10.1074/jbc.M400393200. PubMed PMID: 14734554.

30. Zharkov DO, Golan G, Gilboa R, Fernandes AS, Gerchman SE, Kycia JH, Rieger RA, Grollman AP, Shoham G. Structural analysis of an Escherichia coli endonuclease VIII covalent reaction intermediate. EMBO J. 2002;21(4):789-800. Epub 2002/02/16. doi: 10.1093/emboj/21.4.789. PubMed PMID: 11847126; PMCID: PMC125349.

31. Golan G, Zharkov DO, Feinberg H, Fernandes AS, Zaika EI, Kycia JH, Grollman AP, Shoham G. Structure of the uncomplexed DNA repair enzyme endonuclease VIII indicates significant interdomain flexibility. Nucleic Acids Res. 2005;33(15):5006-16. Epub 2005/09/08. doi: 10.1093/nar/gki796. PubMed PMID: 16145054; PMCID: PMC1199562.

32. Liu M, Doublié S, Wallace SS. Neil3, the final frontier for the DNA glycosylases that recognize oxidative damage. Mutat Res. 2013;743-744:4-11. Epub 2013/01/01. doi: 10.1016/j.mrfmmm.2012.12.003. PubMed PMID: 23274422; PMCID: PMC3657305.

33. Imamura K, Wallace SS, Doublié S. Structural characterization of a viral NEIL1 ortholog unliganded and bound to abasic site-containing DNA. J Biol Chem. 2009;284(38):26174-83. Epub 2009/07/25. doi: 10.1074/jbc.M109.021907. PubMed PMID: 19625256; PMCID: PMC2758016.

34. Liu M, Imamura K, Averill AM, Wallace SS, Doublié S. Structural characterization of a mouse ortholog of human NEIL3 with a marked preference for single-stranded DNA. Structure. 2013;21(2):247-56. Epub 2013/01/15. doi: 10.1016/j.str.2012.12.008. PubMed PMID: 23313161; PMCID: PMC3856655.

35. Imamura K, Averill A, Wallace SS, Doublié S. Structural characterization of viral ortholog of human DNA glycosylase NEIL1 bound to thymine glycol or 5-hydroxyuracil-containing DNA. J Biol Chem. 2012;287(6):4288-98. Epub 2011/12/16. doi: 10.1074/jbc.M111.315309. PubMed PMID: 22170059; PMCID: PMC3281731.

36. Zhu C, Lu L, Zhang J, Yue Z, Song J, Zong S, Liu M, Stovicek O, Gao YQ, Yi C. Tautomerization-dependent recognition and excision of oxidation damage in base-excision DNA repair. Proc Natl Acad Sci U S A. 2016;113(28):7792-7. Epub 2016/06/30. doi: 10.1073/pnas.1604591113. PubMed PMID: 27354518; PMCID: PMC4948311.

37. Campalans A, Marsin S, Nakabeppu Y, O’Connor T R, Boiteux S, Radicella JP. XRCC1 interactions with multiple DNA glycosylases: a model for its recruitment to base excision repair. DNA Repair (Amst). 2005;4(7):826-35. Epub 2005/06/02. doi: 10.1016/j.dnarep.2005.04.014. PubMed PMID: 15927541.

38. Sharma N, Chakravarthy S, Longley MJ, Copeland WC, Prakash A. The C-terminal tail of the NEIL1 DNA glycosylase interacts with the human mitochondrial single-stranded DNA binding protein. DNA Repair (Amst). 2018;65:11-9. Epub 2018/03/10. doi: 10.1016/j.dnarep.2018.02.012. PubMed PMID: 29522991; PMCID: PMC5911420.

39. Dunn AR, Kad NM, Nelson SR, Warshaw DM, Wallace SS. Single Qdot-labeled glycosylase molecules use a wedge amino acid to probe for lesions while scanning along DNA. Nucleic Acids Res. 2011;39(17):7487-98. Epub 2011/06/15. doi: 10.1093/nar/gkr459. PubMed PMID: 21666255; PMCID: PMC3177204.

40. Kladova OA, Grin IR, Fedorova OS, Kuznetsov NA, Zharkov DO. Conformational Dynamics of Damage Processing by Human DNA Glycosylase NEIL1. J Mol Biol. 2019;431(6):1098-112. Epub 2019/02/05. doi: 10.1016/j.jmb.2019.01.030. PubMed PMID: 30716333.

41. Kuznetsov NA, Koval VV, Zharkov DO, Fedorova OS. Conformational dynamics of the interaction of Escherichia coli endonuclease VIII with DNA substrates. DNA Repair (Amst). 2012;11(11):884-91. Epub 2012/09/25. doi: 10.1016/j.dnarep.2012.08.004. PubMed PMID: 23000248.

42. Alexeeva M, Guragain P, Tesfahun AN, Tomkuviene M, Arshad A, Gerasimaite R, Ruksenaite A, Urbanaviciute G, Bjoras M, Laerdahl JK, Klungland A, Klimasauskas S, Bjelland S. Excision of the doubly methylated base N(4),5-dimethylcytosine from DNA by Escherichia coli Nei and Fpg proteins. Philos Trans R Soc Lond B Biol Sci. 2018;373(1748). Epub 2018/04/25 doi: 10.1098/rstb.2017.0337. PubMed PMID: 29685966; PMCID: PMC5915725.

43. Kafer GR, Li X, Horii T, Suetake I, Tajima S, Hatada I, Carlton PM. 5-Hydroxymethylcytosine Marks Sites of DNA Damage and Promotes Genome Stability. Cell Rep. 2016;14(6):1283-92. Epub 2016/02/09. doi: 10.1016/j.celrep.2016.01.035. PubMed PMID: 26854228.

44. Shen B, Chapman JH, Custance MF, Tricola GM, Jones CE, Furano AV. Perturbation of base excision repair sensitizes breast cancer cells to APOBEC3 deaminase-mediated mutations. Elife. 2020;9. Epub 2020/01/07. doi: 10.7554/eLife.51605. PubMed PMID: 31904337; PMCID: PMC6961979.

45. Makasheva KA, Endutkin AV, Zharkov DO. Requirements for DNA bubble structure for efficient cleavage by helix-two-turn-helix DNA glycosylases. Mutagenesis. 2020;35(1):119-28. Epub 2019/12/01. doi: 10.1093/mutage/gez047. PubMed PMID: 31784740.

46. Zhao X, Krishnamurthy N, Burrows CJ, David SS. Mutation versus repair: NEIL1 removal of hydantoin lesions in single-stranded, bulge, bubble, and duplex DNA contexts. Biochemistry. 2010;49(8):1658-66. Epub 2010/01/27. doi: 10.1021/bi901852q. PubMed PMID: 20099873; PMCID: PMC2872175.

47. Liu M, Bandaru V, Bond JP, Jaruga P, Zhao X, Christov PP, Burrows CJ, Rizzo CJ, Dizdaroglu M, Wallace SS. The mouse ortholog of NEIL3 is a functional DNA glycosylase in vitro and in vivo. Proc Natl Acad Sci U S A. 2010;107(11):4925-30. Epub 2010/02/27. doi: 10.1073/pnas.0908307107. PubMed PMID: 20185759; PMCID: PMC2841873.

48. Winn MD, Ballard CC, Cowtan KD, Dodson EJ, Emsley P, Evans PR, Keegan RM, Krissinel EB, Leslie AG, McCoy A, McNicholas SJ, Murshudov GN, Pannu NS, Potterton EA, Powell HR, Read RJ, Vagin A, Wilson KS. Overview of the CCP4 suite and current developments. Acta Crystallogr D Biol Crystallogr. 2011;67(Pt 4):235-42. Epub 2011/04/05. doi: 10.1107/S0907444910045749. PubMed PMID: 21460441; PMCID: PMC3069738.

49. Pei J, Kim BH, Grishin NV. PROMALS3D: a tool for multiple protein sequence and structure alignments. Nucleic Acids Res. 2008;36(7):2295-300. Epub 2008/02/22. doi: 10.1093/nar/gkn072. PubMed PMID: 18287115; PMCID: PMC2367709.

50. Prakash A, Eckenroth BE, Averill AM, Imamura K, Wallace SS, Doublié S. Structural investigation of a viral ortholog of human NEIL2/3 DNA glycosylases. DNA Repair (Amst). 2013;12(12):1062-71. Epub 2013/10/15. doi: 10.1016/j.dnarep.2013.09.004. PubMed PMID: 24120312; PMCID: PMC3856876.

51. Fromme JC, Verdine GL. Structural insights into lesion recognition and repair by the bacterial 8-oxoguanine DNA glycosylase MutM. Nat Struct Biol. 2002;9(7):544-52. Epub 2002/06/11. doi: 10.1038/nsb809. PubMed PMID: 12055620.

