## Supplemental Infomation for "Unique structural features of mammalian NEIL2 DNA glycosylase prime its activity for diverse DNA substrates and environments"

Running Title: Structure of a NEIL2 DNA glycosylase

*To whom correspondence should be addressed: BEE

Funding sources: This work was supported by NIH grant P01 CA098993 awarded to S.D. B.E.E. is supported by NIH grant R50 CA233185.

**Supplemental Table 1.** Anomalous density peaks and isomorphous difference peaks (relative to native) used for phase determination and verification of MdoNEIL2. Shown are the locations of heavy atom sites and associated peak heights; only peaks greater than 3 sigma are shown.

| Residue | Anomalous WT SeMet | L-M SeMet | WT Iodide | WT Au | WT Pt | Isomorphous WT vs. WT SeMet | L-M SeMet vs. WT SeMet | WT vs. Iodide | WT vs. Au | WT vs. Pt |
|---|---|---|---|---|---|---|---|---|---|---|
| M36 | 10.7 | 11.1 | --- | --- | 4.8 | 6.7 | 4.8 | --- | --- | 4.1 |
| M39 | 8.4 | 6.0 | --- | --- | 7.1 | 3.6 | --- | --- | --- | 7.0 |
| M301 | 15.0 | 11.8 | --- | --- | --- | 5.2 | 4.0 | --- | --- | --- |
| L7M | --- | 9.0 | --- | --- | --- | --- | 5.8 | --- | --- | --- |
| L210M | --- | 6.2 | --- | --- | --- | --- | 3.8 | --- | --- | --- |
| L284M | --- | 6.8 | --- | --- | --- | --- | 4.3 | --- | --- | --- |
| C183 | --- | --- | --- | 14.8 | --- | --- | --- | --- | 10.3 | 5.0 |
| R168 | --- | --- | 9.0 | --- | --- | --- | --- | 7.0 | --- | --- |
| C254 | --- | --- |  | 8.9 | 5.0 | --- | --- | --- | 7.3 | --- |
| Zinc | 4.8 | 4.5 | 3.9 | 7.3 | 7.5 | 4.8 | --- | --- | --- | 6.7 |

**Supplemental Figure 1. Comparison of the two molecules of MdoNEIL2 in the crystallographic asymmetric unit.** A.) Shown are the 2 molecules in the asymmetric unit of MdoNEIL2 with chain A colored from N-terminus (blue) to C-terminus (red) and the non-crystallographic two-fold symmetry mate, chain B, shown in grey. B.) Shown is the heavy atom phased and density modified map from AutoSHARP (contoured at 1σ; blue) for the core of the N-terminus and the residual $f_o - f_c$ map (green at +3σ) along a non-crystallographic two-fold axis in the proximity of the insertion (aa 65-131) unique to NEIL2. Also shown are the anomalous difference Fourier peaks at M36 and M39 for each chain contoured at 3σ for the WT selenomethionine dataset (orange). C.) Shown is a least-squares superposition of the two chains with D.) and E.) showing the B-factor representation for chains A and B, respectively. F.) Shown is the orientation of the non-crystallographic two-fold axis relative to the C-cell edge (Green and cyan) vs. the initial phasing solution at higher symmetry assuming the two-fold to be crystallographic (grey).

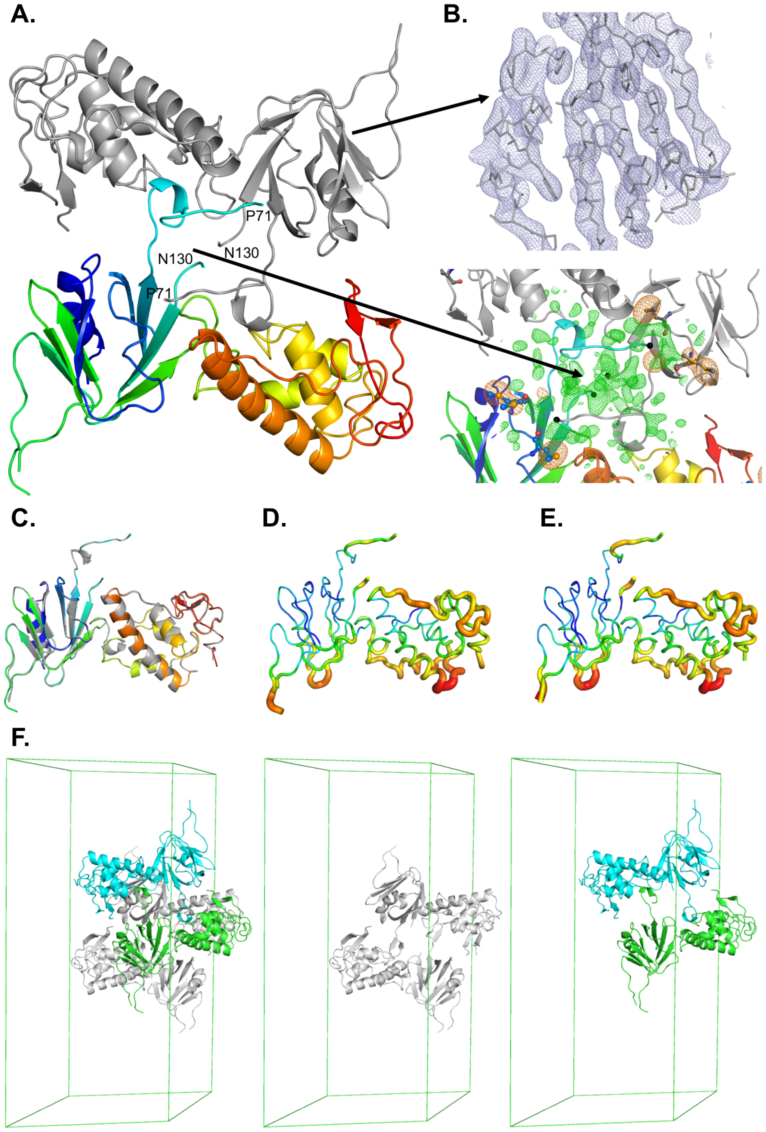

**Supplemental Figure 2. Comparison of full length MdoNEIL2 and MdoNEIL2cut crystals shows isomorphism and expected minor differences.** A.) Shown is the $f_o - f_o$ isomorphous difference Fourier map contoured at +/-3σ (green and red, respectively) between MdoNEIL2 full-length selenomethionine data and the MdoNEIL2cut construct ($f_{o(Se)} - f_{o(cut)}$) at the non-crystallographic two-fold axis containing the large insert. The green map along the non- crystallographic axis indicates the longer connection between P71 and N130 for MdoNEIL2 while the red mesh indicates the shorter connection for MdoNEIL2cut. MdoNEIL2cut retains up to S67 and the lack of difference density in the N-terminal direction indicates the local structure is retained for the construct. B.) Shown is the same map at the Se site (M301) along one of the NCS two-folds. The green density is reflective of the higher electron content of the Se atom. C.) Shown is the isomorphous difference Fourier map contoured at +/-3σ (green and red, respectively) between MdoNEIL2 gold derivative and the MdoNEIL2cut construct ($f_{o(Au)} - f_{o(cut)}$) at C183 on the left and C254 on the right.

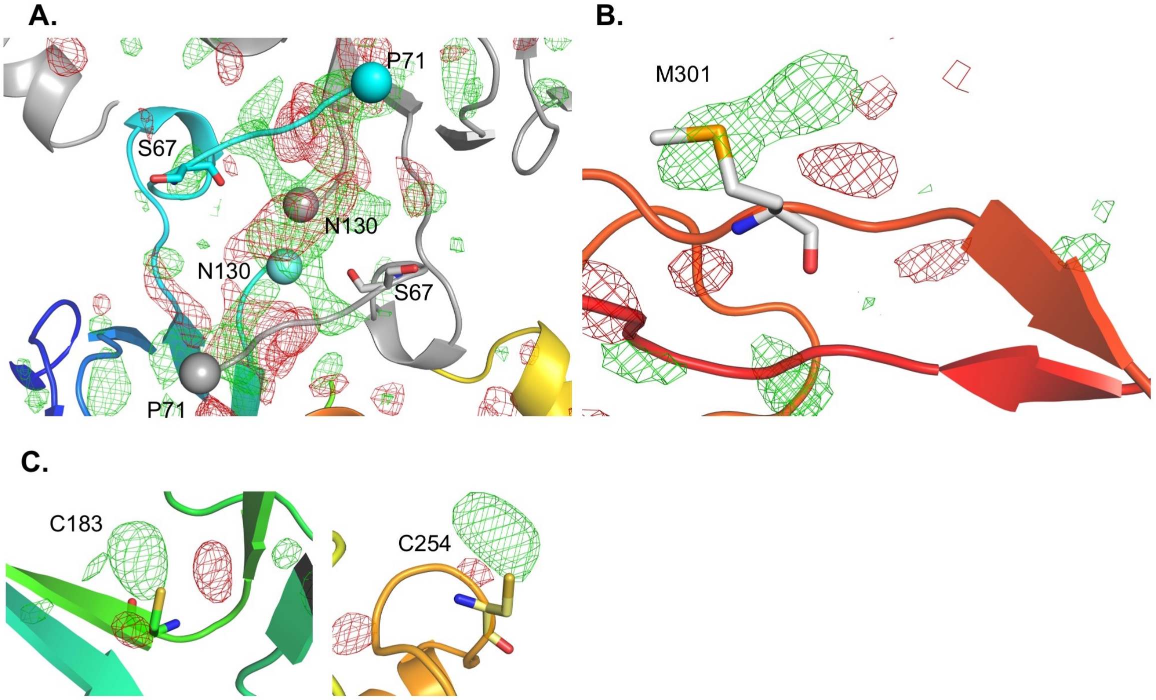

**Supplemental Figure 3. Biochemical characterization of NEIL2.** A.) Glycosylase activity comparison of NEIL2 variants under single-turnover conditions for $^{32}$P-labelled single stranded abasic site containing DNA substrate run on urea-PAGE. Assays were performed using 25 nM DNA substrate and enzyme concentration titrated from 0 to 800 nM. B.) Shown is the activity titration for single stranded DNA substrates. C). Shown is the activity titration for double stranded DNA substrates.

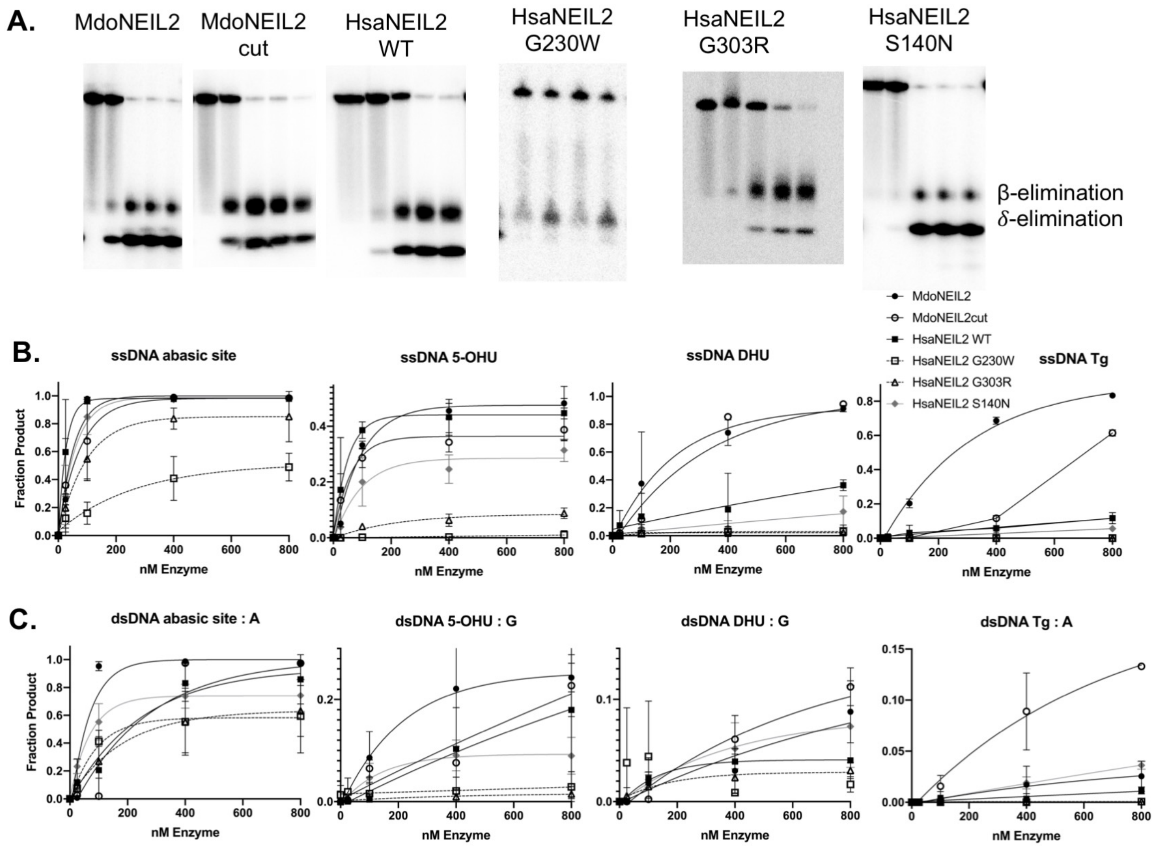

**Supplemental Figure 4.** Structure-based sequence alignment of the Fpg/Nei family glycosylases produced using PROMALS3D(1). The sequences used in the alignment are MvNei1 (PDB ID code: 3A46)(2), HsaNEIL1 (1TDH)(3), MmuNEIL3 (3W0F)(4), MvNei2/3 (4MB7)(5), LlaFpg (1L1T)(6), and EcoNei (2EA0).

```
                         PE
MvNei1              1   MPEGPEVALTADILEKYF-KGKTLEYI-DFI-SGRYSKSEPE---------------------GYDDFIA   46
HsaNEIL1            1   MPEGPELHLASQFVNEAC-RALVFGGCVEKSSVSRNP----------------------------EVPFE   41
MdoNEIL2            1   MPEGPSLRKFHQLVAPF--VGQLV-VTVGGN--SKKI----------------------------NPNM-   36
HsaNEIL2            1   MPEGPLVRKFHHLVSPF--VGQQV-VKTGGS--SKKL----------------------------QPAS-   36
MmuNEIL3            1   MVEGPGCTLNGEKIRARVLPGQAV-TGVRGTALQ-SLLGPAMSPAASLADVATSAAPMNAKDSGWKLLRL   67
MvNei2/3            1   MVEAPRIRITYEKIRHT--KNHRI-VSISGP--SYK-----------------------------RMNVD   35
LlaFpg              1   MPELPEVETIRRTLLPLI-VGKTI-EDVRIF-WPNIIRHPRD---------------------SEAFAAR   46
EcoNei              1   MPEGPEIRRAADNLEAAI-KGKPLTDV-WFA-FPQ----------------------------LKTYQSQ   38
Consensus aa:           .sEtP.lp.h.p.l..h...tp.l.........sp..............................p....
Consensus ss:                hhhhhhhhhhhhh    ee eeee                                  hhhhhh

                                     K                           Large Insert NEIL2
MvNei1             47   NLPLKVSNVDTKGKFLWFELFDPND-----KSN-------------------------------------   74
HsaNEIL1           42   SSAYR-ISASARGKELRLILSPLPGA--QPQQE-------------------------------------   71
MdoNEIL2           37   LEMLRLQDSQVHGKNLYLNFGLTEDLGLPESFLLPKHLQKKVRLPKEKSDHKLETTSRLDGQEVPGSSLA  106
HsaNEIL2           37   LQSLWLQDTQVHGKKLFLRFDLDEEMGPPGSSPTPEPPQKEVQKEGAADPKQVGEPSGQKT----LDGSS  102
MmuNEIL3           68   FNGYVYSGVETLGKELFMYFG-------------------------------------------------   88
MvNei2/3           36   LIDYIIRKWWFAGKYIYLMLISS-----------------------------------------------   58
LlaFpg             47   MIGQTVRGLERRGKFLKFLLD-------------------------------------------------   67
EcoNei             39   LIGQHVTHVETRGKALLTHFSN------------------------------------------------   60
Consensus aa:           ..sb.lp.hph.GK.Lbh.h.................................................
Consensus ss:           h   eeeeeeee  eeeeee

                                                     Void Fill  Small Insert NEIL2
MvNei1             75   -------------------------KWYIWNTFGLTGMWSLFEA--------------KYTRAVLSFDN-  104
HsaNEIL1           72   -------------------------PLALVFRFGMSGSFQLVPREE----------LPRHAHLRFYTAPP  106
MdoNEIL2          107   IKALELGEEEKETVMPWWLNTSQNSGLWLCFHFGLFGSVRASELSRATKANKRGDWKDPIPRLVLHFAK-  175
HsaNEIL2          103   RSAELVPQGEDDSEYLERDAPAGDAGRWLRVSFGLFGSVWVNDFSRAKKANKRGDWRDPSPRLVLHFGG-  171
MmuNEIL3           89   -------------------------PRALRIHFGMKGSILINPREG-------ENRAGASPALAVQLTR-  125
MvNei2/3           59   ---------------------NKPTYVIRTHMMMHGRILVGNQDS-----------PTKRAFMIIQLDN-   95
LlaFpg             68   -------------------------RDALISHLRMEGRYAVASALEP---------LEPHTHVVFCFTD-  102
EcoNei             61   -------------------------DLTLYSHNQLYGVWRVVDTGEEPQ-------TTRVLRVKLQTAD-   97
Consensus aa:           .........................hl..ph.h.G.h.ls...p................s.h.h.hs..
Consensus ss:                                     eeeeee   eeeeeee                 eeeeeee

                       Orphaned Base  Wedge             Interdomain Linker Region
MvNei1            105   --ELMAYFSDMRNFGTFKFSNS-----EKELKRKLN-ELGPDFLK-NDDIDISKIKK-------------  152
HsaNEIL1          107   GPRLALCFVDIRRFGRWDLG-----------GKWQP-GRGPCVLQEYQQFRESVLRNLAD----------  154
MdoNEIL2          176   ---GFLAFY----NCRIYWCL--------------GPTV--KPTSDILS--EEFDRRQALEALK------  214
HsaNEIL2          172   --GGFLAFY----NCQLSWSS--------------SPVV--TPTCDILS--EKFHRGQALEALG------  211
MmuNEIL3          126   ---DLICFY----DSSVELRN----SVE------SQQRVRVMEELDICS--PKFSFSRAESEVK------  171
MvNei2/3           96   --DIVLRWY----RSQITLLDPNCLAEIKTNYTICTTRQAIMDSIKLMKYDLSNNRFDYNLFQSHLKNGI  159
LlaFpg            103   --GSELRYRDVRKFGTMHVYAKEE----ADRRPPLA-ELGPEPL--SPAFSPAVLAERAVK---------  154
EcoNei             98   --KTILLY---SASDIEMLRPEQ---------LTTHPFLQRVGPDVLD--PNLTPEVVKERLLS------  145
Consensus aa:           ....hl.@........s.p.b........................hhp...php...hbp.l....
Consensus ss:              eeeeee       eeeeee                hh hhhh  h            hhhhhhhhhh


                             H2TH Motif  Lesion Backbone
MvNei1            153   ---YKQPIVALLMDQKKIGSGLGNYLVAEILYRAKIDPHKLGSNLTDQEIENLWYWIKYET---------  210
HsaNEIL1          155   -KAFDRPICEALLDQRFF-NGIGNYLRAEILYRLKIPPFEKARSVLEALQQHRPSPELTLSQKIRTKLQN  222
MdoNEIL2          215   ---QASPVSYTLLDQRYF-AGLGNIIKNEVLYLARIHPLSLGSCLTPLNLE-------------------  261
HsaNEIL2          212   ---QAQPVCYTLLDQRYF-SGLGNIIKNEALYRAGIHPLSLGSVLSASRRE-------------------  258
MmuNEIL3          172   -KQGDRMLCDVLLDQRVL-PGVGNIIKNEALFDSGLHPAVKVCQLSDKQAC-------------------  219
MvNei2/3          160   NIHSSEIITDFLLDQEYF-PGVGNILQQEALYDCKILPLKKVQDIDEPMFD-------------------  209
LlaFpg            155   ---TKRSVKALLLDQTVV-AGFGNIYVDESLFRAGILPGRPAASLSSKEI--------------------  200
EcoNei            146   PRFRNRQFAGLLLDQAFL-AGLGNYLRVEILWQVGLTGNHKAKDLNAAQLDALAHALLEIP---------  205
Consensus aa:           ....pp.ls.hLLDQphh.sGlGNhl.sEhL@ph.l.P.pbs.sls..b.p...................
Consensus ss:                  hhhhhh          hhhhhhhhhhh   hhhhh    hhhhh

                                                                     Lesion Backbone
MvNei1            211   -----------K---LAYDSN----HIGYMVNLENESSKIGRKNYHPNIHPTEKEFDFLVYRKKKDPNGN  262
HsaNEIL1          223   PDLLELCHSVPK---EVVQLGGRGYGSESGEEDF-----------------AAFRAWLRCYGM-------  265
MdoNEIL2          262   -SLLDHVVSFSV---GWLQKKLE--G-------------------------KPL--HHLIYQK--EQCP-  295
HsaNEIL2          259   -VLVDHVVEFST---AWLQGKFQ--G-------------------------RPQ--HTQVYQK--EQCP-  292
MmuNEIL3          220   -HLVKMTRDFSI---LFYRCCKA--G-------------------------SAISKHCKVYKR--PNCD-  255
MvNei2/3          210   -CLCNSLKKIIDLLYESYKFRESGKE-------------------------FGP--ILRIYRK--SLCP-  248
LlaFpg            201   ERLHEEMVATIG---EAVMKG----GSTV-RTYVNT-----------QGEAGTFQHHLYVYGRQGNPCK-  250
EcoNei            206   -----------R---FSYATR----GQVDEN------------------KHHGALFRFKVFHRDGEPCE-  238
Consensus aa:           ...............b.h..................................s....h.lY.+..p.C..
Consensus ss:            hhhhhhhhhhh   hhhhh                                    eeee

                                    Lesion Backbone
MvNei1            263   KVIADKIIGSG-----KNKRTTYWAPAIQKLE--------------------------------------  289
HsaNEIL1          266   ----PGMSSLQD----RHGRTIWFQGDPGP-LAPKGRKSRKKKSKATQLSPEDRVEDALPPSKAPSRTRR  326
MdoNEIL2          296   --AGHQVMKDSFGPPGSFQRLTWWCPHCQPKAEEKVEVTQEQL---------------------------  336
HsaNEIL2          293   --AGHQVMKEAFGPEDGLQRLTWWCPQCQPQLSEEPEQCQFS----------------------------  332
MmuNEIL3          256   -QCHSKITVCRFGE---NSRMTYFCPHCQ-----------------------------------------  280
MvNei2/3          249   --LGH---KTIRKKIGLRNRMTTWCPVCQL----------------------------------------  273
LlaFpg            251   -RCGTPIEKTV-----VAGRGTHYCPRCQR----------------------------------------  274
EcoNei            239   -RCGSIIEKTT-----LSSRPFYWCPGCQH----------------------------------------  262
Consensus aa:           ..htp.l............R.h@@tP.hQ.........................................
Consensus ss:                eeeeeeeee  e  eeeee
```

**Supplemental Figure 5. Evaluation of *E. coli* endonuclease VIII and MdoNEIL2 conformations by SAXS.** Shown is the conformational analysis using MultiFoXS(7) for EcoNei (A) and MdoNEIL2cut (B) with the experimental scattering curve along with the fits to 1-5 states (conformations) displayed. For both proteins, the $X^2$ minima could be achieved by inclusion of three states for each model. C.) Shape analysis using Dammif(8) for EcoNei with the model on the left showing the top 10 selected envelopes and the green surface representation showing the average produced by Damaver(9). Also shown is a fit to the unliganded crystal structure (PDB ID 1Q3B), with the projection likely representing the flexible loop (aa 213-223) in the C-terminal domain. D.) Similar analysis of MdoNEIL2 with Dammif generated two populations of envelopes with 5 (left) showing some agreement and the other 5 (right) appearing random.

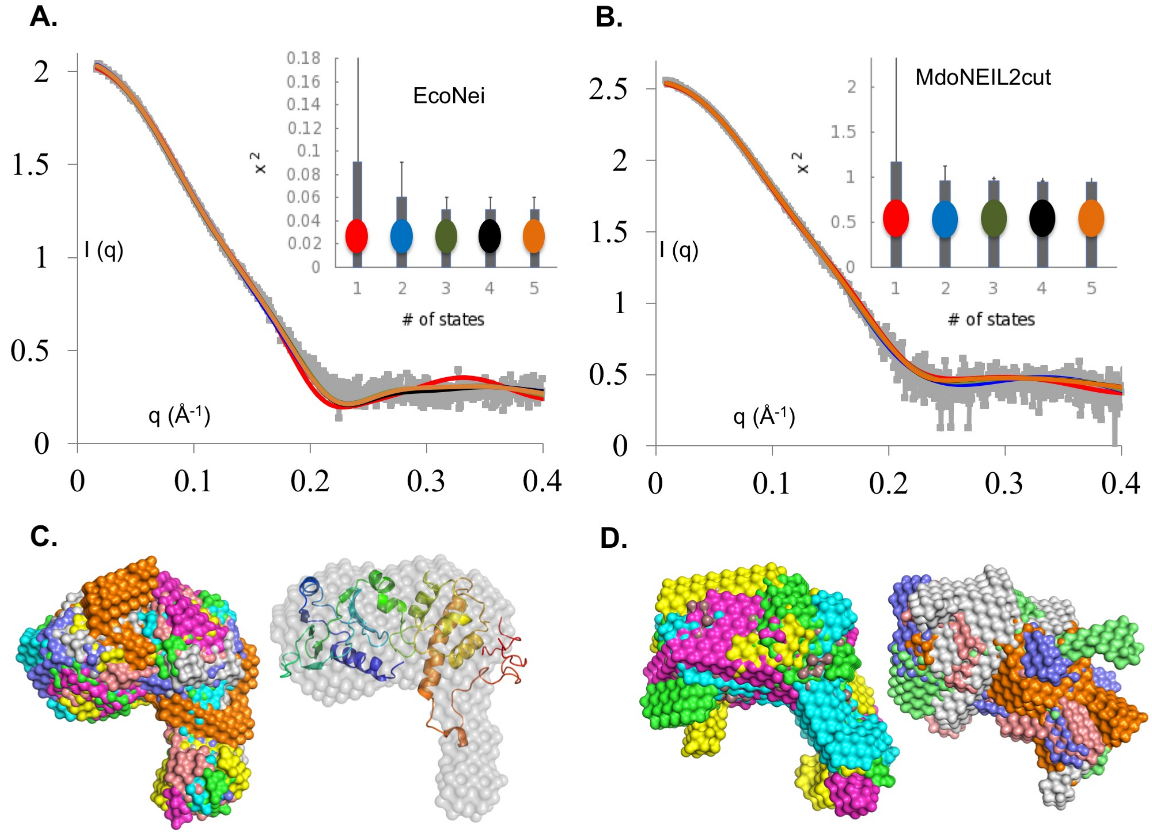

### REFERENCES


1. Pei J, Kim BH, Grishin NV. PROMALS3D: a tool for multiple protein sequence and structure alignments. Nucleic Acids Res. 2008;36(7):2295-300. Epub 2008/02/22. doi: 10.1093/nar/gkn072. PubMed PMID: 18287115; PMCID: PMC2367709.
2. Imamura K, Wallace SS, Doublié S. Structural characterization of a viral NEIL1 ortholog unliganded and bound to abasic site-containing DNA. J Biol Chem. 2009;284(38):26174-83. Epub 2009/07/25. doi: 10.1074/jbc.M109.021907. PubMed PMID: 19625256; PMCID: PMC2758016.
3. Doublié S, Bandaru V, Bond JP, Wallace SS. The crystal structure of human endonuclease VIII-like 1 (NEIL1) reveals a zincless finger motif required for glycosylase activity. Proc Natl Acad Sci U S A. 2004;101(28):10284-9. Epub 2004/07/03. doi: 10.1073/pnas.0402051101. PubMed PMID: 15232006; PMCID: PMC478564.
4. Liu M, Imamura K, Averill AM, Wallace SS, Doublié S. Structural characterization of a mouse ortholog of human NEIL3 with a marked preference for single-stranded DNA. Structure. 2013;21(2):247-56. Epub 2013/01/15. doi: 10.1016/j.str.2012.12.008. PubMed PMID: 23313161; PMCID: PMC3856655.
5. Prakash A, Eckenroth BE, Averill AM, Imamura K, Wallace SS, Doublié S. Structural investigation of a viral ortholog of human NEIL2/3 DNA glycosylases. DNA Repair (Amst). 2013;12(12):1062-71. Epub 2013/10/15. doi: 10.1016/j.dnarep.2013.09.004. PubMed PMID: 24120312; PMCID: PMC3856876.
6. Fromme JC, Verdine GL. Structural insights into lesion recognition and repair by the bacterial 8-oxoguanine DNA glycosylase MutM. Nat Struct Biol. 2002;9(7):544-52. Epub 2002/06/11. doi: 10.1038/nsb809. PubMed PMID: 12055620.
7. Schneidman-Duhovny D, Hammel M, Tainer JA, Sali A. FoXS, FoXSDock and MultiFoXS: Single-state and multi-state structural modeling of proteins and their complexes based on SAXS profiles. Nucleic Acids Res. 2016;44(W1):W424-9. Epub 2016/05/07. doi: 10.1093/nar/gkw389. PubMed PMID: 27151198; PMCID: PMC4987932.
8. Franke D, Svergun DI. DAMMIF, a program for rapid ab-initio shape determination in small-angle scattering. Journal of Applied Crystallography. 2009;42:342-6. doi: 10.1107/S0021889809000338. PubMed PMID: WOS:000264292800030.
9. Volkov VV, Svergun DI. Uniqueness of ab initio shape determination in small-angle scattering. Journal of Applied Crystallography. 2003;36:860-4. doi: 10.1107/S0021889803000268. PubMed PMID: WOS:000182284400105.
